# Magnetic torque-driven living microrobots for enhanced tumor infiltration

**DOI:** 10.1101/2022.01.03.473989

**Authors:** T. Gwisai, N. Mirkhani, M. G. Christiansen, T.T. Nguyen, V. Ling, S. Schuerle

**Author notes:** Corresponding author, Address: ETH Zurich, Vladimir-Prelog-Weg 1-5/10, CH-8093 Zürich, Switzerland.

## Abstract

Bacterial microrobots combining self-propulsion and magnetic guidance are increasingly recognized as promising drug delivery vehicles for targeted cancer therapy. Thus far, control strategies have either relied on poorly scalable magnetic field gradients or employed directing magnetic fields with propulsive forces limited by the bacterial motor. Here, we present a magnetic torque-driven actuation scheme based on rotating magnetic fields to wirelessly control *Magnetospirillum magneticum* AMB-1 bearing versatile liposomal cargo. We observed a 4-fold increase in conjugate translocation across a model of the vascular endothelium and found that the primary mechanism driving this increased transport is torque-driven surface exploration at the cell interface. Using spheroids as a 3D tumor model, fluorescently labeled bacteria colonized their core regions with up to 21-fold higher signal in samples exposed to rotating magnetic fields. In addition to enhanced transport, we demonstrated the suitability of this magnetic stimulus for simultaneous actuation and inductive detection of AMB-1. Finally, we demonstrated that RMF significantly enhances AMB-1 tumor accumulation *in vivo* following systemic intravenous administration in mice. Our findings suggest that scalable magnetic torque-driven control strategies can be leveraged advantageously with biohybrid microrobots.

**One-Sentence Summary:** Magnetic torque-driven motion enhances infiltration of living microrobots across physiological barriers both *in vitro* and *in vivo*.

## INTRODUCTION

Bacteria that self-propel and preferentially colonize tumors are poised to become next generation living microrobots for cancer therapy (*1, 2*). Unlike conventional chemotherapeutics reliant on passive diffusion for transport, these living microrobots possess attributes that enable them to actively overcome challenges posed by the pathological characteristics of the tumor environment (*3*–*5*). Tumor-targeting bacteria are particularly appealing because of their capacity to autonomously navigate through the body, transport a wide range of payloads, and modulate intratumoral inflammatory responses (*6*–*9*). To date, strains of *Salmonella, Mycobacterium*, and *Clostridium* have been tested in animal models and progressed to clinical trials for bacteria-based cancer therapy (*10*–*13*). Nevertheless, translation of this approach has been hindered by incomplete clinical responses, in part due to insufficient tumor colonization (*14*). Developing control strategies to enhance and accelerate accumulation of bacteria at the target site is essential to limit immune clearance, facilitate robust colonization, and ultimately increase therapeutic efficacy.

Stimuli including chemical gradients, light, electric fields, and magnetic fields have been employed for wireless control of bacteria (*15*–*18*). Of these, magnetic fields are especially promising for medical use due to their precise spatiotemporal targeting, minimally-invasive deep tissue penetration, and well-established clinical safety (*19*). Recently, innately magnetic strains of bacteria acting as steerable therapeutic microrobots have been manipulated with external magnetic fields (*20*). In their native aquatic habitats, magnetotactic bacteria (MTB) biomineralize magnetite or greigite nanocrystals and use magnetically assisted aerotaxis to migrate to regions of low oxygen concentration. *In vivo*, strains of MTB carrying payloads have been shown to preferentially proliferate in deoxygenated regions of tumors following peritumoral injection in the presence of a directional magnetic field (DMF) or a magnetic field gradient (*21, 22*). While these studies illustrate the promise of MTB as drug delivery agents, their potential for clinical translation is curbed by reliance on peritumoral administration, which is suitable only for easily accessible tumors, as well as limitations associated with the magnetic stimuli they employ. Using DMF inherently relies on MTB self-propulsion, which limits tissue penetration capability (*23*). Drawing MTB toward target sites with static field gradients also has fundamental shortcomings, especially in the context of deep targets, since magnetic field gradients rapidly diminish with increasing distance from their source (*24, 25*).

In this work, we establish a hybrid control strategy that harnesses magnetic torque-driven motion followed by autonomous taxis-based locomotion to enhance the infiltration of *Magnetospirillum magneticum* AMB-1 as a carrier for covalently-coupled liposomes (MTB-LP) (Fig. 1). Unlike some forms of magnetic stimulus, uniform rotating magnetic fields (RMF), like those employed here, can be generated at clinically relevant scales for deep sites within the body. Our previous work used MTB exposed to RMF to generate local fluid convection to enhance the transport of non-magnetic nanoparticles across collagen matrices in microfluidic models (*26*). Here, we instead show how RMF can be used to significantly enhance the infiltration of living microrobots themselves across robust tissue barriers. Using the Caco-2 Transwell system, we establish that RMF increases MTB translocation across this characteristically impermeable barrier, compared to DMF or the absence of magnetic actuation. We also demonstrate the suitability of RMF for simultaneous actuation and inductive detection, which could be exploited for closed-loop operation and real-time monitoring. We then assess MTB-LP infiltration in the presence of RMF using model tissue barriers. By studying extravasation with computational models and *in vitro*, we find that the main mechanism driving the enhancement of translocation is increased surface exploration resulting from torque-driven translational motion at the cell interface. We assess the spatiotemporal characteristics of magnetically enhanced MTB-LP infiltration and colonization in a 3D tumor model and study the effects of RMF on MTB flank tumor accumulation *in vivo* following systemic intravenous administration. We conclude that the MTB-LP platform, when combined with an RMF actuation scheme, is a versatile biohybrid system that could improve targeting and colonization of therapeutic bacteria as living microrobots in tumors.

**Fig. 1:**
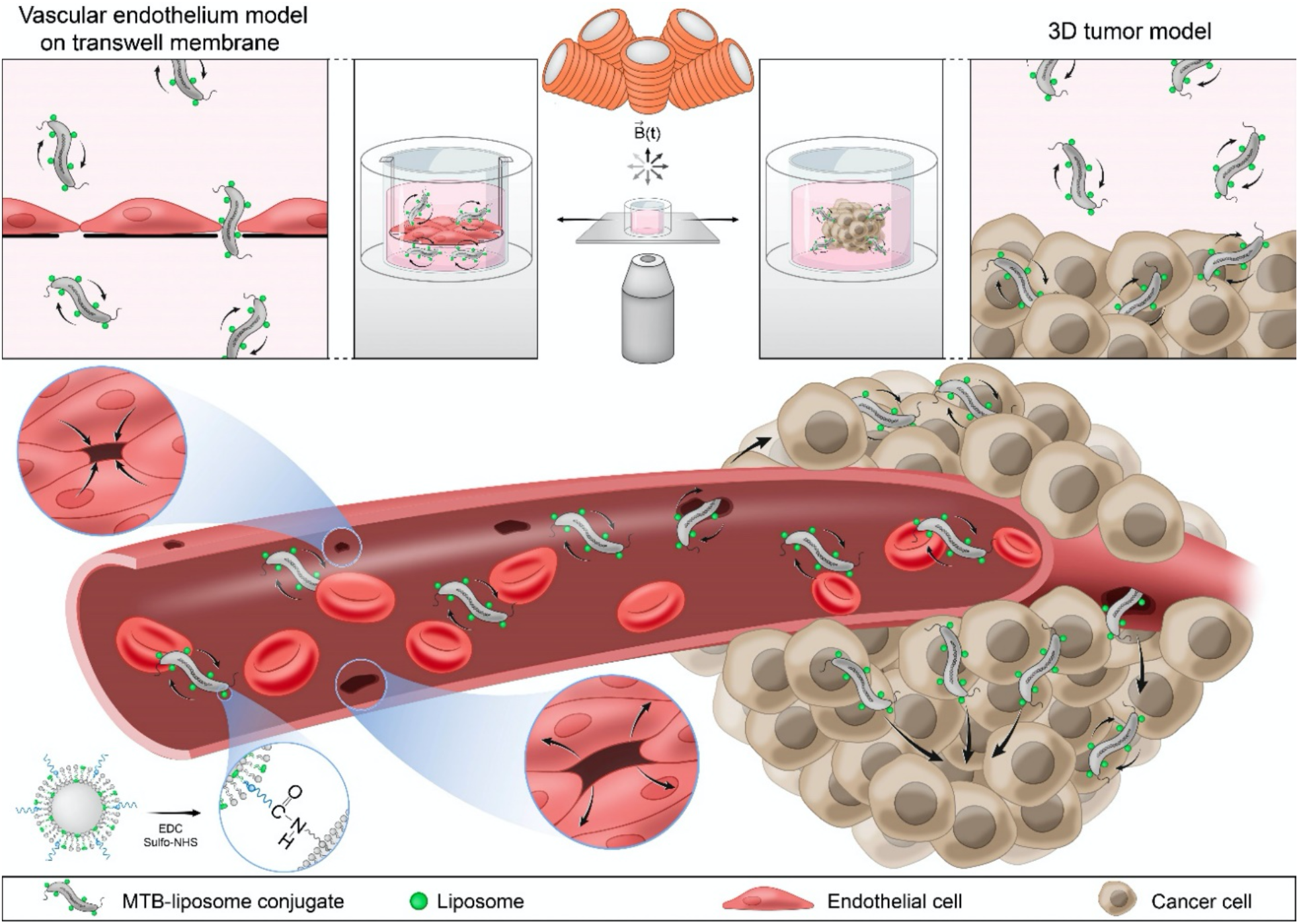
Conceptual overview of magnetically enhanced MTB-liposome conjugate transport. Integrated magnetic manipulation and imaging setup used in experiments to analyze the transport of MTB-liposome conjugates in a vascular endothelium model and a 3D tumor model (bottom). Samples are placed between the objective lens and electromagnetic coils which generate a uniform rotating magnetic field, and MTB-liposome conjugates rotate individually in response to the applied field. Schematic illustrating magnetically induced transport of MTB-liposome conjugates through a blood vessel and into a tumor (bottom).

## RESULTS

### Rotating magnetic fields enhance translocation and enable inductive detection of MTB

A range of strategies have been employed to manipulate MTB as living microrobots using external magnetic fields (Fig. 2A). In the presence of DMF, the motion of the bacteria is dependent on the propulsive force (*F*_*P*_) generated by their flagella and the fluidic drag force (*F*_*D*_), which are equal and opposite when traveling at constant velocity. An estimate of the propulsive force is given by *F*_*P*_ = *γv*, where *γ* is the linear drag coefficient and *v* is the linear velocity. For a velocity range of 19 - 49 µm/s (*27, 28*), the propulsive force is estimated to be on the order of 0.1 pN. Static field gradients have also been employed to pull MTB towards a target site. While this approach has shown efficacy in small animal models, an impracticable gradient of more than 130 T/m would be required to produce a comparable force on the same magnetic moment.

**Fig. 2:**
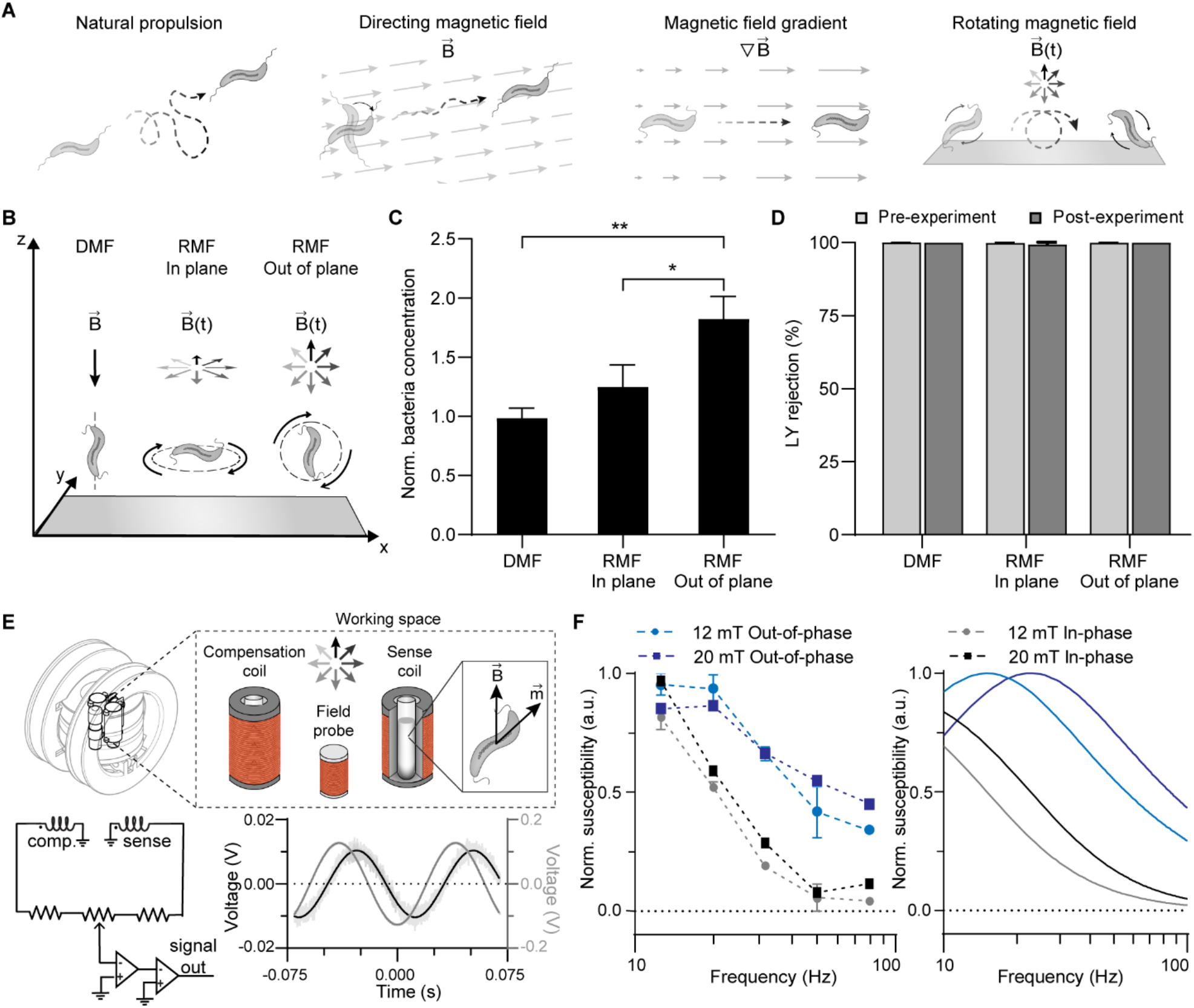
Rotating magnetic fields enhance translocation and enable inductive detection of MTB. (**A**) Schematic illustrating various magnetic actuation strategies employed to manipulate magnetotactic bacteria. (**B**) Schematic depicting the reference coordinate system used in all actuation experiments, and orientation of AMB-1 after exposure to directional static magnetic fields (DMF) and in-plane and out-of-plane rotating magnetic fields (RMF). (**C**) Comparison of normalized basolateral bacteria concentrations following 1 hour of exposure to either DMF, in-plane rotation or out-of-plane rotation, both at 14 Hz. In these experiments only, a magnetic force force generated by spatial variation of the magnitude of the RMF or DMF (1 T/m) was applied along -z. Bacteria concentration was normalized to respective unactuated controls. (*n* = 3; mean ± SD; \**P* < 0.05 and \*\**P* < 0.01, ANOVA). (**D**) Pre- and post-experimental Lucifer yellow (LY) rejection values for DMF and in-plane and out-of-plane RMF (*n* = 3; mean ± SD). (**E**) Cancellation and amplification scheme for RMF magnetometer. A typical signal detected from the MTB (measured at 12 mT at 12.6 Hz) and signal recorded from the in-phase field probe are shown. (**F**) Experimental in-phase and out-of-phase susceptibility (left; *n* = 3; mean ± SD) and analytical susceptibility data (right).

In contrast to DMF, AMB-1 self-propulsion is overridden when subjected to RMF. Viscous drag from the surrounding fluid creates a phase lag between the external field and magnetic moment of the MTB, giving rise to a magnetic torque that is exerted on the bacteria. The opposing hydrodynamic torque (*τ*_*H*_) that defines the steady state lag angle between the MTB and magnetic field is governed by *τ*_*H*_ *= α ω*, where *α* is the rotational drag coefficient and the angular frequency (*ω*) is equal to the frequency of magnetic actuation under synchronous rotation. An estimate of the force generated by this torque-based motion is given by 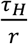, where *r* is the distance from the axis of rotation, which is half the bacterial length. For angular frequencies in the range of 10 – 25 Hz and distance *r* of 0.9 µm (fig. S1), the force generated is on the order of 1 pN. These estimates show that, on average, magnetic torque-driven motion generates forces that are an order of magnitude higher than those of MTB self-propulsion.

We sought to examine whether the higher forces and torque-based motion produced by RMF can enhance the infiltration of MTB across biological barriers. We first characterized our actuation strategy under stringent conditions, employing a low field amplitude and a characteristically impermeable barrier. Caco-2 cells cultured on Transwell membranes have been used extensively for the reconstitution of differentiated enterocyte-like monolayers to evaluate oral drug absorption and the translocation of various strains of bacteria (*29*–*31*).

MTB were added to the upper chamber of a Transwell insert with an established Caco-2 monolayer and placed in a PDMS well containing media. The bacterial concentration in the PDMS well was determined using a hemocytometer after one hour of magnetic actuation at 12 mT and 14 Hz. MTB were rotated in-plane (parallel) and out-of-plane (orthogonal) with respect to the cell monolayer and compared to exposure to DMF alone which was applied along -z (Fig. 2B). Under the stringent conditions of these initial cell monolayer experiments only, a magnetic force generated by spatial variation of the magnitude of the RMF or DMF (1 T/m) was applied along -z to support translocation. The RMF maintained its plane of rotation at all points.

Out-of-plane rotation was found to significantly increase MTB penetration compared to both in-plane RMF and DMF (Fig. 2C). Out-of-plane actuation for frequencies between 1-20 Hz consistently resulted in higher translocation than controls without magnetic exposure (fig. S2). Given these findings, all subsequent experiments were performed using out-of-plane RMF.

To verify that MTB actuation does not affect monolayer integrity, the Lucifer yellow (LY) rejection assay was performed before and after each experiment. This fluorescent molecule passively diffuses through the cell monolayer by paracellular transport only and, thus, is unable to cross Caco-2 monolayers when tight junctions are well established. LY rejection values remained above 99% in all tested conditions, indicating that no damage occurred to the monolayer (Fig. 2D and fig. S3).

Having observed enhanced MTB translocation using RMF, we sought to investigate the use of rotating magnetic fields for detection of MTB response. Unlike DMF or gradient fields, the time-varying nature of RMF can be exploited for simultaneous actuation and monitoring of MTB, enabling the adjustment of actuation parameters during operation for improved localization and infiltration.

In a proof-of-concept experiment, inductive detection of MTB during actuation was performed using a small scale, custom-built RMF magnetometer consisting of two pairs of nested Helmholtz coils (Fig 2E, fig. S4). Phase shifted sinusoidal currents were applied to each set of coils to produce a circularly rotating magnetic field. When exposed to RMF, the overall magnetic moment of the MTB varies in time by rotating, allowing the measurement of the resulting voltage induced in surrounding loops of wire. Inductive field probes for each pair of coils were placed in the central workspace along with symmetric sense and compensation coils that served to isolate inductive signal from the magnetization of the MTB. Following fine tuning of their cancellation via a potentiometer, the resulting signal was amplified and acquired. The magnetization of the MTB was phase shifted relative to the rotating field, reflecting the phase lag in the magnetic response of the bacteria.

Using a cosine function fitted to the in-phase field probe, integration was used to separate the acquired signal into in-phase and out-of-phase components of the susceptibility of the sample (Fig. 2F). The experimental data was compared to analytical data to assess whether the detected signal generated by the MTB exhibited predicted characteristics (Supplementary Materials). Out-of-phase susceptibility at 12 mT and 20 mT peaked at 12.6 Hz and 20 Hz respectively, compared to 14 Hz and 24 Hz in the analytical data. The underlying trends for both the in-phase and out-of-phase susceptibility were as predicted by our models, confirming that we were able to detect MTB under actuation with rotating magnetic fields. Overall, these studies illustrate the promise of RMF for both detection and remote actuation of MTB for enhanced transport.

### Elucidating the role of torque-driven motion on translocation using computational modelling

Given the prominent effect of RMF on MTB transport across robust cell barriers, we sought to understand the main mechanism driving enhanced translocation using a computational model in COMSOL Multiphysics. Since the endothelium is the first biological barrier encountered following intravenous administration, transport across a 2D endothelial cell monolayer was modeled. Torque exerted on the bacteria upon exposure to RMF results in forces that are applied on the cell surface, as well as the generation of substantial translational motion. Our estimates above show that, on average, magnetic torque-driven motion generates forces (∼ 1 pN) that are an order of magnitude higher than those of MTB self-propulsion, yet lower than those required to break bonds formed by vascular endothelial-cadherin (VE-cadherin) (*32*), an endothelial cell-specific adhesion molecule. This suggests that the application of forces directly on an endothelial monolayer cannot be the chief mechanism responsible for enhanced translocation. As such, we sought to examine the influence of translational motion derived from torque-based actuation on bacterial translocation.

An endothelial cell monolayer was modeled with adjacent cells forming a sealed barrier between upper and lower compartments. The cells were modeled as hyperelastic materials with a shear modulus of 1 kPa (*33, 34*) and dimensions adopted from Arefi *et al*. (*34*). Recent work has shown that dynamic mechanical processes within the endothelium result in gap formations that are independent of the influence of migrating cancer or immune cells (*35, 36*). Using this as a basis, stochastic opening of cell-cell contacts was incorporated into the model to account for the active mechanics of the endothelium. For each simulation, the gap lifetime was set to 160 s (35) and a set of random parameters were generated to determine the gap size, which was within the range of 1.5 to 2.5 μm (34). The overall simulation time was selected to encompass opening incidences for all gaps. Considering the relative stiffness of gram-negative bacteria compared to endothelial cells, MTB were modeled as rigid ellipsoids possessing a rigid dipole moment along the long axis. TEM images and multisizer data were used to determine the approximate size of a single bacterium, which were estimated to be 0.45 and 1.8 μm for the short and long axis respectively (fig. S1) (*37*). Hydrodynamic interactions were modeled as linear and rotational viscous damping acting on the rigid body.

To characterize the permeability of our modeled monolayer, simulations of passive diffusion of liposomes with a diameter of approximately 200 nm were performed and 5.9% of the liposomes diffused into the lower compartment (Fig. 3A, Movie S1). Having established the model, we proceeded to compare MTB transport under DMF and RMF (Fig. 3B, Movie S2 and S3). The velocity profiles generated for the bacteria under RMF exhibited the well-studied characteristics of surface walkers under low Reynolds number flows (59–61). The higher mass density of MTB with respect to the surrounding liquid gives rise to a terminal velocity of the bacteria which results in an offset in the y component. When traveling along the monolayer, there is discontinuous contact with the surface and the contributions from the x and y components of the velocity vary depending on whether the MTB is traveling downhill, uphill or is translocating. As anticipated, the contribution from the x component is minimal as the MTB passes through an opening, reflecting the lowered contact with the cell surface.

**Fig. 3:**
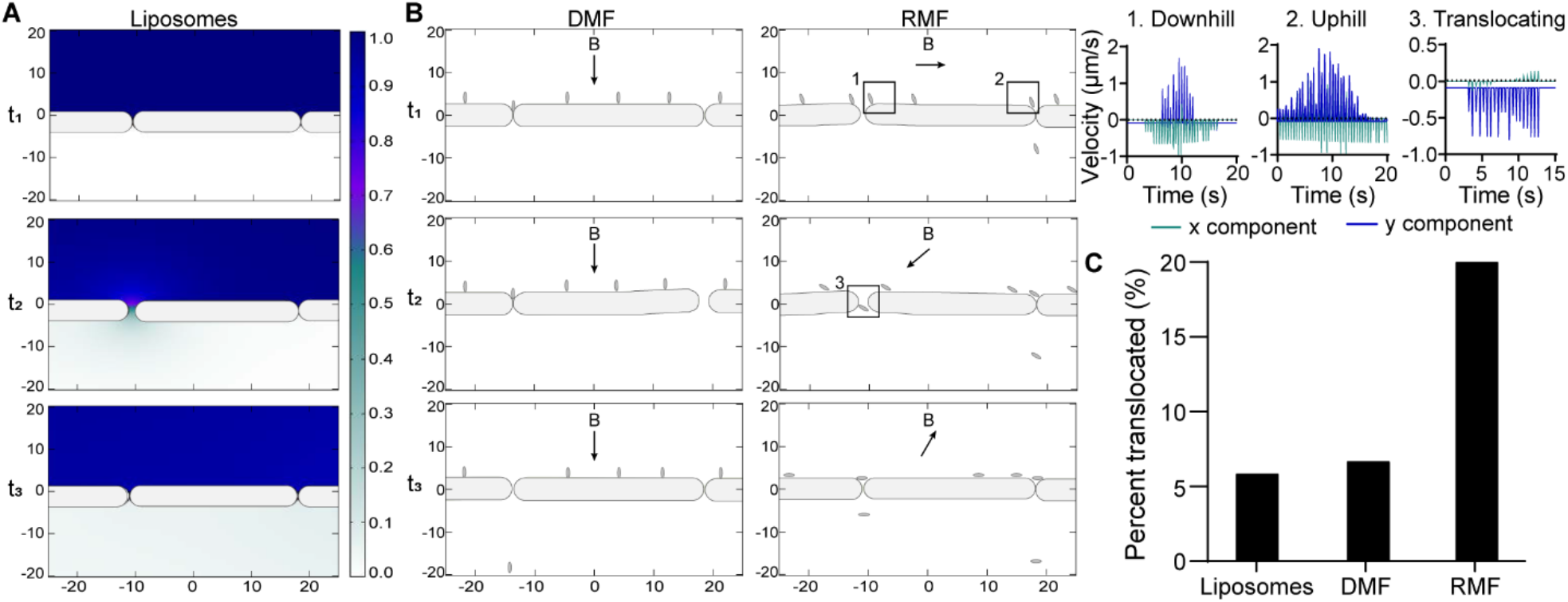
Computational modeling of MTB-LP transport across endothelial monolayers. (**A**) Computational simulations of liposome diffusion across a modelled endothelial monolayer. Cells were modeled as hyperelastic materials and Stochastic opening of cell-cell contacts was incorporated into the model to account for spontaneous gap formation. (**B**) Simulations MTB transport under DMF and RMF across a modelled endothelial monolayer. MTB were modeled as rigid ellipsoids possessing a magnetic dipole moment. Velocity profiles show contributions of x and y components of MTB traveling downhill (1), uphill (2), and translocating through a cell gap (3). (**C**) Plot of the average amount of liposome diffusion and MTB translocation under DMF and RMF for 3 simulations relative to the starting concentrations.

Our model showed that only 6.6% of MTB exposed to DMF crossed the barrier, compared to 20% of MTB exposed to RMF (Fig. 3C). Under the static conditions of DMF, the bacteria only passed through the monolayer when initially located in close proximity to a cell-cell junction. In contrast, MTB under RMF translated along the monolayer which enabled the bacteria to explore the monolayer surface and pass through any gaps that formed between the cells. Overall, these findings strongly suggest that enhanced surface exploration resulting from torque-based translational motion is the dominant mechanism facilitating increased translocation of MTB.

### Rotating magnetic fields increase extravasation of MTB-liposome conjugates

We proceeded to investigate translocation in an *in vitro* system suited to model vascular extravasation. Human microvascular endothelium (HMEC-1) monolayers cultured on Transwell inserts were used to mimic the endothelial barriers that would be encountered following intravenous delivery. To verify that intact monolayers were formed when the cells were cultured on permeable membranes, immunostaining for VE-cadherin was performed (Fig. 4A). Confocal imaging showed the presence of intact cell-cell contacts and the formation of a uniform monolayer after 2 days. In an initial experiment, passive diffusion of unconjugated liposomes across the monolayer was measured (fig. S5). From the fluorescence intensity values, it was found that only 0.24% of the liposomes were able to cross into the basolateral chamber via passive diffusion after 1 hour (Fig. 4B).

**Fig. 4:**
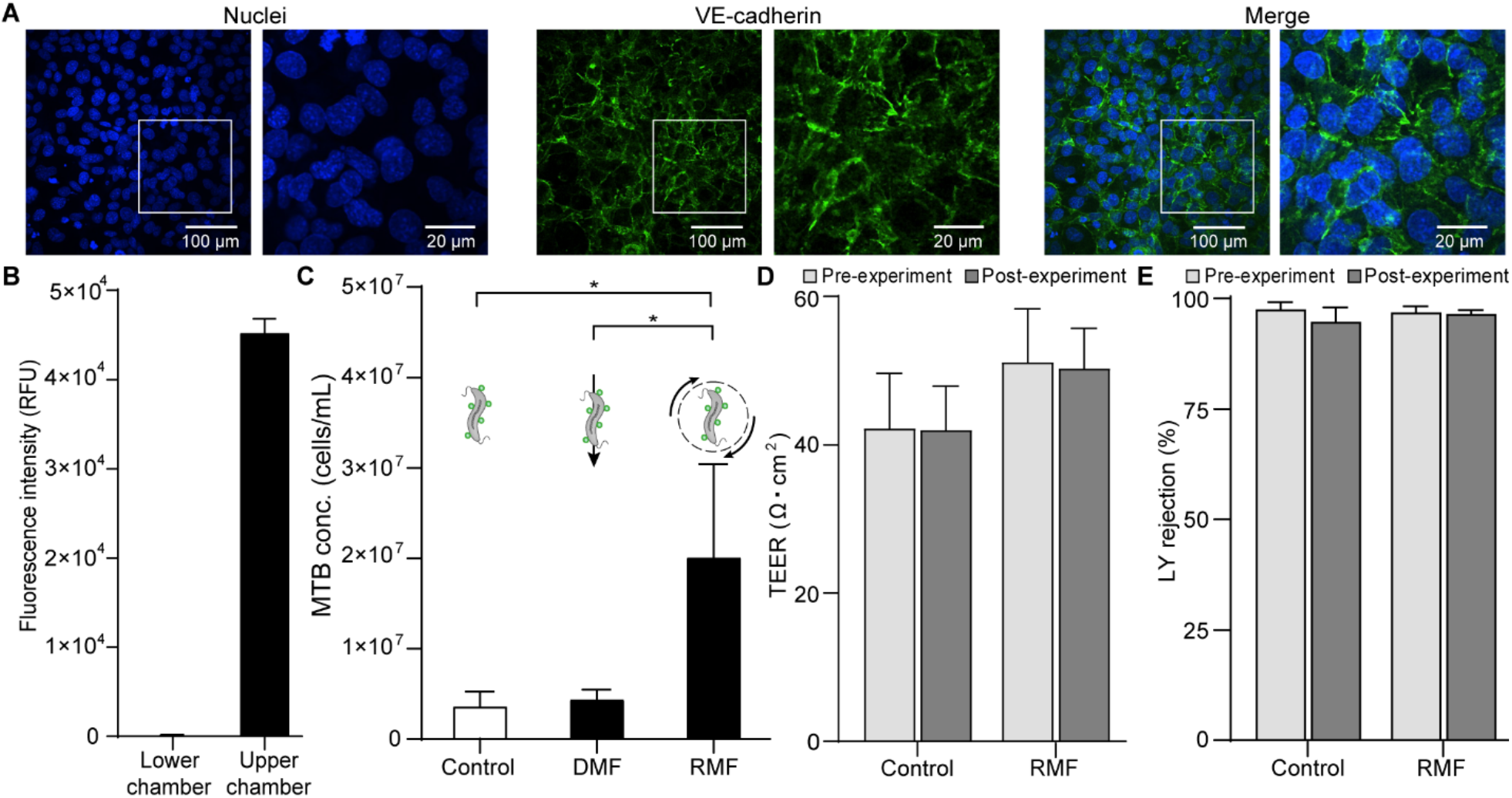
Increasing translocation across endothelial monolayers. (**A**) Representative confocal images of immunostaining for VE-cadherin (green) on HMEC-1 cells cultured on a Transwell membrane. Cell nuclei were stained using Hoechst 33342 (blue). (**B**) Evaluation of the lower and upper compartment fluorescence intensity measurements following 1 hour of passive diffusion of liposomes across HMEC-1 monolayers (n = 3; mean ± SD). (**C**) Comparison of lower compartment MTB-LP concentrations after 1 hour for unactuated controls and exposure to either DMF (12 mT) and out-of-plane RMF (20 mT and 24 Hz) (n = 3; mean ± SD; *P < 0.05, ANOVA). No gradients were applied in these experiments. (**D**) Pre- and post-experimental transendothelial electrical resistance (TEER) measurements for unactuated controls and out-of-plane RMF (*n* = 3; mean ± SD). (**E**) Pre- and post-experimental Lucifer yellow (LY) rejection values for unactuated controls and out-of-plane RMF (*n* = 3; mean ± SD).

We then investigated the translocation of MTB-liposome conjugates across the monolayer. The MTB–liposome complex (MTB–LP) combines the adaptability of traditional polymeric drug carriers and the functionality of magnetic based platforms to produce a living biohybrid vehicle for targeted drug delivery. Bioconjugation of fluorescently labelled liposomes to the MTB cell surface was achieved through carbodiimide mediated amidation (fig. S6, Table S1). Conjugates were added to the upper chamber of Transwell inserts and the concentration in the lower chamber was determined after one hour of actuation. To increase the available magnetic torque, which is a function of magnetic moment and magnetic field strength, the applied field magnitude was increased to 20 mT. At this field strength, the step out frequency of our MTB strain of interest was found to be 24 Hz (*37*), in agreement with the magnetometer data (Fig. 2F).

MTB-LP were actuated out-of-plane relative to the monolayer for 1 hour, and this was compared to exposure to DMF, as well as unactuated controls (Fig. 4C). Out-of-plane rotation led to 4.6-fold higher MTB-LP translocation compared to DMF. Exposure to DMF resulted in MTB-LP concentrations that were comparable to unactuated controls, as was the case in the Caco-2 monolayer experiments (Fig. 2C). Transendothelial electrical resistance (TEER) measurements and the LY rejection assay were performed to verify that MTB actuation does not affect monolayer integrity (Fig. 4D and 4E). Both TEER and LY rejection values before and after actuation were comparable, demonstrating that disruption of the monolayer did not occur due to exposure to magnetic actuation.

### Rotating magnetic fields improve MTB transport and colonization of 3D tumor spheroids

Having demonstrated enhanced translocation across endothelial monolayers, we next sought to examine the effect of RMF on MTB-LP tumor penetration in an *in vitro* tumor model using human breast adenocarcinoma (MCF-7) spheroids. Tumor spheroids can effectively recapitulate the mass transport properties and complex architecture of avascular tumor tissue and have been used to study bacteria-based cancer therapies (*38*–*40*). High levels of E-cadherin expression in MCF-7 cells facilitate the formation of strong cell-cell adhesions particularly in 3D spheroid cultures, making them an ideal model for difficult-to-penetrate tumor masses (*41*). MCF-7 cells transitioned from loose aggregates to highly compact 3D spheroids after 3 days of culture. The spheroids were transferred to PDMS wells, into which conjugates were added. MTB suspensions were stained with a proliferation dye prior to liposome conjugation to visualize daughter cells over the course of the experiment. After one hour of actuation at 20 mT and 24 Hz, spheroids were washed thoroughly and incubated for up to 120 hours. Given that comparable results for unactuated controls and DMF were comparable (Fig. 2C, Fig. 4C), as well as lack of a clearly defined preferential direction for steering MTB in an inherently 3D model, RMF was compared to unactuated controls for all subsequent experiments.

Confocal images of live tumors were captured at increments of 10 μm from the bottom of the spheroids to visualize the distribution of DiO-labelled liposomes after 24 h of incubation (Fig. 5A). MTB-LP conjugates were able to propel into deep regions of avascular spheroids following exposure to RMF. The density of the conjugate clusters detected in the actuated samples increases with increasing depth. This finding was confirmed by evaluating the mean intensity of each section and compiling a fluorescence distribution for the z-plane up to a depth of 100 μm (Fig. 5B). The fluorescence intensity values were consistently higher for actuated samples compared to controls and peak fluorescence for actuated samples occurred at 80 µm. Decreased signal after 80 µm could be caused by limitations in optical sectioning of deep tissue regions. Given that MTB-LP were mostly located in the center of actuated samples, the high cell density in this region results in tissue opacity, light scattering, and limited light penetration which restricts visualization of deep tissue sections by confocal microscopy (*42*). Nevertheless, it was still evident that a significantly higher number of conjugates penetrated the spheroids in RMF-exposed samples.

**Fig. 5:**
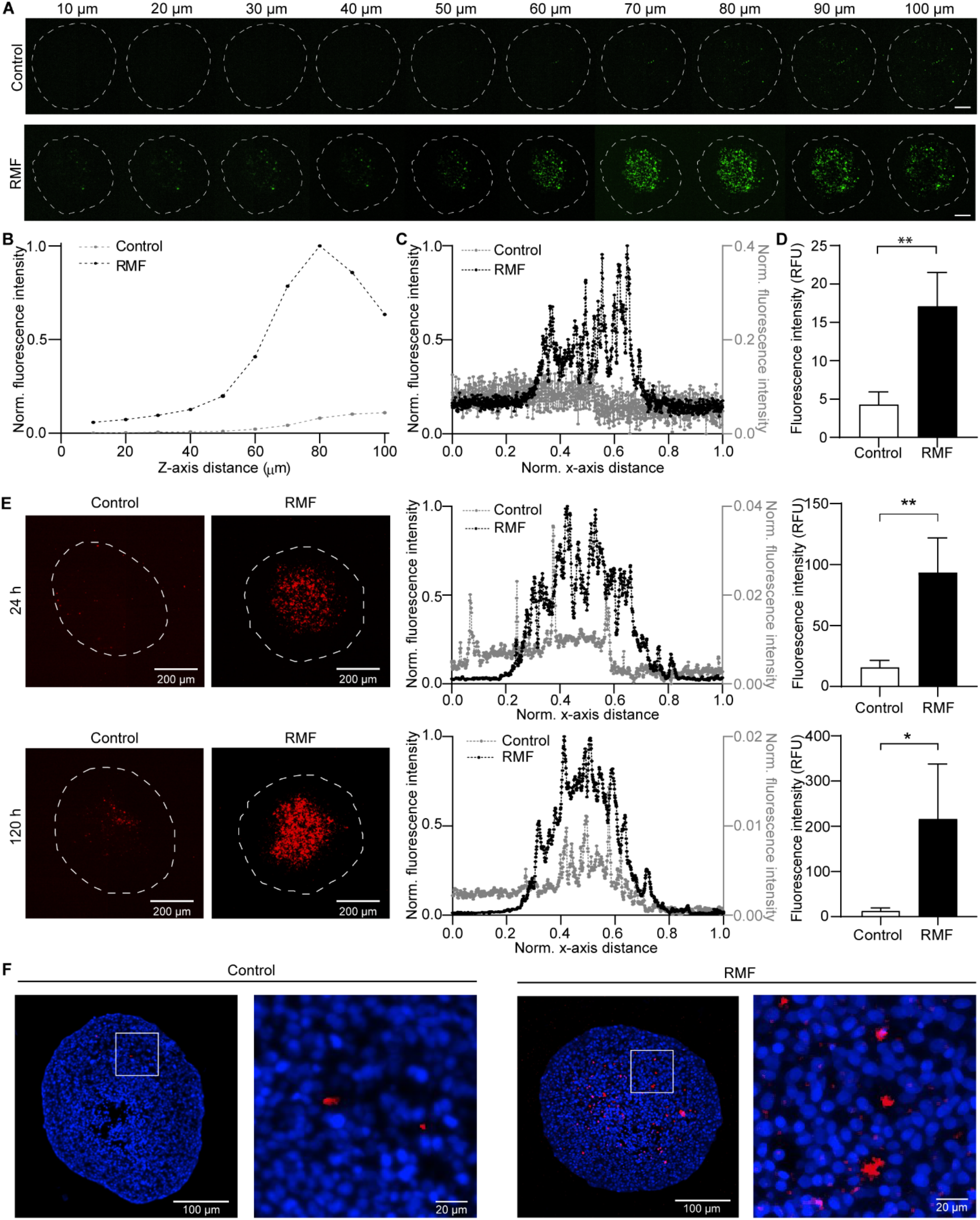
Intratumoral transport of MTB conjugates in 3D tumor spheroids. (**A**) Representative confocal images of live MCF-7 spheroids following 1 hour exposure to RMF (20 mT and 24 Hz), thorough washing, and 24 hours incubation. Control refers to unactuated samples. Images were captured at 10 μm increments from the bottom of the spheroids and show the localization of DiO-labelled liposomes (green). Outline depicts the shape of the spheroid at its largest circumference. Scale bar = 100 μm. (**B**) Plot of mean intensity for each section at 10 μm intervals from the bottom of the spheroids along the z-axis up to a depth of 100 μm. (**C**) Representative fluorescence intensity distribution of the 80 µm section. Values were normalized to overall minimum and maximum fluorescence intensity values. Spheroid diameter was normalized along the x-axis. (**D**) Summation of mean intensity values for consecutive Z-plane images up to a depth of 100 μm (*n* = 3; mean ± SD; \*\**P* < 0.01, Student’s t-test). (**E**) Representative z-projections of MTB stained with a far-red proliferative dye in live MCF-7 spheroids following 1 hour exposure to RMF (20 mT and 24 Hz), thorough washing, and incubation for up to 120 h without actuation. Control refers to unactuated samples. Images were captured at 24 and 120 h. Outline depicts the shape of the spheroid at its largest circumference. Representative normalized fluorescence intensity distribution of actuated samples and controls at 24 and 120 h (center). Values were normalized to overall minimum and maximum fluorescence intensity values. Spheroid diameter was normalized along x-axis. Image-based quantification of fluorescence intensity values from z-projections at 24 and 120 h (right; *n* = 3; mean ± SD; \**P* < 0.05, \*\**P* < 0.01, Student’s t-test). (**F**) Representative images of 5 μm histology sections for MCF-7 spheroids 144 h after actuation.

The intensity distribution profiles for the 80 µm sections show distinct differences in the distribution of MTB-LP in actuated samples compared to controls (Fig. 5C). Control samples showed a more uniform distribution of the fluorescence signal. In contrast, the overall fluorescence intensity values were higher in actuated samples and the fluorescence signal was highest in the central region of the spheroid. A summation of mean fluorescence intensity for each section up to a depth of 100 μm showed that the overall fluorescence intensity, and thus the amount of conjugates delivered in the actuated samples is 4-fold higher than in the controls (Fig. 5D).

Having characterized the efficacy of conjugate delivery using RMF, we were interested in investigating MTB-LP tumor colonization over time. Confocal images of live tumors were used to visualize and quantify the distribution of fluorescently labelled bacteria at 24 and 120 hours for RMF-exposed samples and unactuated controls (Fig. 5E). The z-projection images revealed that MTB were still detectable within the spheroids for both actuated samples and controls after 120 hours. Fluorescence intensity profiles of the spheroids were used to study the localization of the MTB in the spheroids. MTB achieved colonization of the core regions of both actuated and unactuated samples. Additionally, there was a narrowing in the intensity profile, indicating an increased density of bacteria at the center of the spheroid from 24 to 120 hours. This enhanced accumulation may be the result of autonomous taxis-based locomotion. Spheroids over 400 μm in diameter can develop a hypoxic core, and preferential accumulation of MTB in this region might result from oxygen-sensing mechanisms that facilitate their navigation towards hypoxic environments (*43*–*45*).

A summation of mean fluorescence intensity for each section up to a depth of 100 μm was used to assess the relative amount of MTB present in the spheroids at various time points. The fluorescence intensity values in spheroids exposed to RMF were 9.9-fold and 21.3-fold higher than in the controls at 24 and 120 hours, respectively. The ability of RMF to enhance transport and colonization of MTB was also investigated using HCT 116 spheroids (fig. S7). As with MCF-7 spheroids, higher MTB colonization in the core regions of the spheroids was achieved in actuated samples after 120 hours. Importantly, over the course of the experiments the bacteria did not replicate uncontrollably, which is likely a result of the relatively low doubling time of MTB compared to other strains of bacteria. This suggests that some of the risks of infectious complications which have hampered the use of some bacterial strains could be of low concern (*10*). To overcome the challenges associated with imaging whole spheroids, histological sectioning of MCF-7 spheroids was performed (Fig. 5F). Sections from the center of the spheroids showed that MTB formed clusters between the cells and that there was higher signal from the MTB in spheroids exposed to RMF compared to controls. Overall, these findings indicate that the combination of magnetic torque-driven motion with taxis-based navigation results in robust tumor colonization.

### Assessment of intratumoral transport of MTB *in vivo*

Motivated by the pronounced effect of RMF on MTB transport *in vitro*, we next sought to test whether our RMF actuation strategy also enhanced bacterial accumulation *in vivo* with a pilot study employing a syngeneic mouse model. BALB/c nude mice bearing subcutaneous MCF-7 tumors in one hind flank received intravenous injections of 1×10^9^ MTB stained with a far-red proliferative dye (Fig. 6A). The anaesthetized mice were placed either in the absence of magnetic actuation (control) or with tumors positioned in the workspace of a magnetic field generator applying a 20 mT RMF at 14 Hz for 1 h. For an applied field magnitude of 20 mT, uniform fields with negligible offset gradients are expected within the workspace of the field generator (Fig. 6B, fig. S8), ensuring that observed effects on the accumulation of bacteria can solely be attributed to the rotational character of the applied field.

**Fig. 6:**
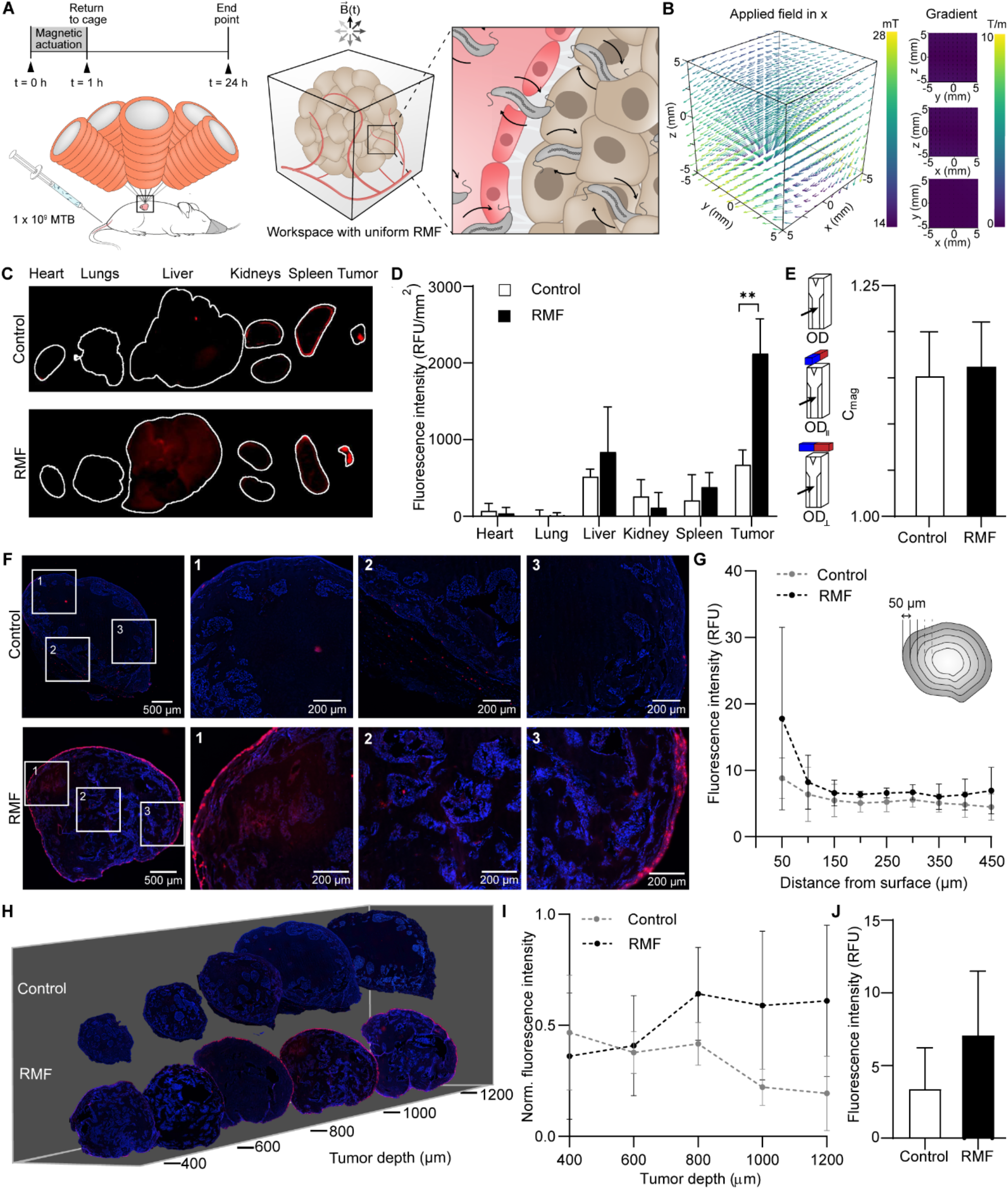
Assessment of intratumoral transport of MTB *in vivo*. (**A**) BALB/c nude mice bearing subcutaneous MCF-7 tumors in one hind flank were intravenously administered with 1 × 10^9^ MTB stained with a far-red proliferative dye. The mice were anaesthetized for 1 h in the absence of exposure to magnetic actuation (control) or were placed on a magnetic field generator with tumors positioned in the workspace (RMF). Mice were returned to the cage for 24 h, after which the tumor and major organs were harvested for further analysis. (**B**) Plots of expected field magnitude and gradients produced in the workspace of the magnetic field generator for an applied field of 20 mT in x. (**C**) Representative *ex vivo* fluorescence intensity images of major organs and tumors 24 h after injection of far-red stained MTB. **(D)** Quantitative biodistribution from harvested organs and tumors (*n* = 3; mean ± SD; \*\**P* < 0.01, Student’s t-test). **(E)** C_mag_ values for homogenized tumors placed in liquid culture for 8 days (*n* = 3 control, *n* = 4 RMF; mean ± SD). **(F)** Representative images of 10 μm histology sections which were sectioned at depth of approximately 1000 μm in the tumour. Cell nuclei were stained using Hoechst 33342 (blue). **(G)** Mean intensity values with increasing distance from the periphery of the tumor sections (n = 3; mean ± SD). **(H)** Representative transverse tumor sections. Cell nuclei were stained using Hoechst 33342 (blue). **(I)** Plot of mean intensity for each traverse tumor section at 200 μm intervals. Values were normalized to overall minimum and maximum fluorescence intensity values. (**J**) Summation of mean intensity values for consecutive traverse tumor sections (*n* = 3; mean ± SD).

Following treatment, mice were returned to the cage for 24 h, after which the tumor and major organs were harvested to assess the distribution of the bacteria using a whole organ fluorescence scanner (Fig. 6C). Fluorescence signal was detected in all tumors suggesting that the bacteria innately accumulate in tumors. Notably, this signal was 3.16-fold higher in mice with tumors exposed to RMF compared to unactuated controls (Fig. 6D). Very low signals were detected in most of the major organs except for the liver, consistent with previous findings 24 h after intravenous administration of AMB-1 (*46*). Bacterial accumulation in the liver is expected to decrease over time, with full clearance by day 6 (*46*).

To assess whether the bacteria detected in the fluorescence scans were still viable, harvested tumors were homogenized and placed into MTB culture media. After 8 days, large, dark pellets that were magnetically responsive were present in all tubes, indicating that the tumor homogenates contained live MTB. C_mag_ measurements were performed to quantify the magnetic properties of the cultures and all samples had values above 1, signifying magnetic responsiveness of the bacterial suspensions (Fig. 6E).

Histological sectioning of the tumors was performed, with sections collected at a depth of approximately 1 mm in the tumor showing more pronounced accumulation at the periphery in both control and actuated samples (Fig. 6F). Mean intensity distributions with increasing distance from the periphery for the tumor sections show that bacteria can be detected throughout the measured region, with higher overall signal from samples exposed to RMF compared to controls (Fig. 6G). This result indicates higher penetration for RMF actuated tumors.

Transverse histology sections were also collected to assess the distribution of MTB with increasing depth within the tumor (Fig. 6H). The mean intensity of each section was used to compile a fluorescence intensity distribution (Fig. 6I). The intensity increases with increasing depth in the first 800 μm of RMF-exposed samples that were analyzed, whereas intensity decreases with depth in the controls. The average fluorescence intensity for consecutive transverse tumor sections showed that the overall fluorescence intensity, and thus the amount of MTB delivered in the actuated samples, is 2.1-fold higher than in the controls (Fig. 6J). These results, combined with the findings from our *in vitro* analysis, demonstrate the potential of magnetic torque-driven control strategies for enhanced tumor accumulation.

## DISCUSSION

The development of strategies to enhance tumor targeting and infiltration are essential for the clinical translation of bacterial cancer therapy which, to date, has been hindered by difficulties in achieving sufficient tumor colonization. Efforts to enhance the accumulation of therapeutic agents using magnetic fields have most often relied on powerful static field gradients to draw drug carriers towards a target site (*47*–*49*). This approach has intrinsic limitations that narrow its potential for clinical translation. Magnetic fields decrease with the cube of the distance from a dipole and their gradients diminish even more rapidly, implying that the length scales associated with many tumors would require sources with impracticable field strengths. In contrast, uniform rotating magnetic fields can be generated at clinically relevant scales for actuation of magnetically-responsive vectors in deep sites.

The dynamic nature of RMF may eventually be exploited for the dual purposes of control and detection of magnetic agents. We established with proof-of-concept experiments that MTB actuated with RMF can be inductively detected, laying a foundation for further development of simultaneous monitoring and closed-loop control strategies. *In vivo* inductive detection of magnetic tracers driven with alternating magnetic fields has already been widely investigated with magnetic particle imaging (MPI) (*50*). The possibility for concurrent imaging and actuation with MPI has also been recognized, combining particle tracking of superparamagnetic iron oxide nanoparticles with cancer hyperthermia *in vivo* (*51*). Gradient fields that select signal from image voxels in MPI have proven challenging to scale up to humans, but recent efforts have demonstrated MPI in deep regions in human brains with moderate resolution (*52*). Our recent work has shown that torque-based actuation, as well as the resulting inductive signals generated by the MTB, should be compatible with selection fields (*37*). One foreseeable challenge to simultaneous inductive monitoring and actuation of MTB with RMF is that frequencies of tens of hertz are used rather than the kilohertz signals generated in MPI, resulting in correspondingly weaker inductive signals. Another is the lack of higher order harmonics contributed to the inductive signal with rotational actuation, which arise in MPI from periodic saturation of the magnetic tracers. Future work will need to identify strategies to overcome these obstacles.

Using *in vitro* models for various biological barriers, we demonstrated that a hybrid control strategy harnessing magnetic torque-driven motion followed by autonomous taxis-based locomotion enhances the transport of living microrobots. MCF-7 spheroids were used to model avascular tumor tissue while the Transwell system was used to generate a model for a typically impermeable barrier using Caco-2 cells, and HMEC-1 cells were used to mimic the vascular endothelium. In both monolayer models, translocation driven by application of an RMF significantly outperformed the DMF and unactuated controls (Fig. 1B and Fig. 3C). Computational modeling showed that torque-driven translational motion, which led to increased cell surface exploration, was the primary mechanism responsible for increased transport across monolayers. Although not the chief mechanism driving MTB-LP transport, shear forces exerted by rotating MTB-LP on the cell surface may still contribute in part to higher transport across monolayers and increased accumulation in spheroids. Moreover, the heterogeneity of MTB populations may allow a small subset of bacteria with especially high magnetic moments to exert sufficient forces on mechanotransducers, such as VE-cadherin and E-cadherin, to trigger responses including transient disruptions of intercellular junctions (*53, 54*). Combined with the translation of the bacteria on the cell surface, these relatively rare disruptions could assist in enhancing local permeability and transport of MTB-LP.

We also showed that the results from the *in vitro* characterization of our actuation strategy can be replicated in the highly complex biological environment *in vivo*. Given the viability of AMB-1 upon intravenous administration, future studies will incorporate longer timepoints to facilitate the study of taxis-based accumulation in tumor cores, as well as bacterial clearance from off-target sites. Notably, we have demonstrated that significant accumulation of MTB can be achieved for tumors exposed to RMF following systemic administration. Previous studies employing magnetic actuation following peritumoral administration have shown considerable bacterial accumulation, but this route of administration is limited to easily accessible tumors, which raises the barrier to practical clinical use.

With a demonstrated ability to overcome robust biological barriers, our actuation strategy is a potentially powerful tool for improving the targeting and efficacy of magnetically manipulable therapeutic agents. The versatility of the MTB-LP platform potentiates the delivery of a wide variety of therapeutic cargo without employing genetic engineering. By merging the benefits of bacteria-mediated delivery with a scalable magnetic torque-driven actuation scheme, our approach enables biohybrid microrobots to help overcome persistent transport barriers and improve cancer treatment.

## MATERIALS AND METHODS

### Materials

Human colorectal adenocarcinoma (Caco-2, ATCC HTB-37), human microvascular endothelium (HMEC-1, ATCC CRL-3243), human breast adenocarcinoma (MCF-7, ATCC HTB-22), human colorectal carcinoma (HCT 116, ATCC CCL-247), *Magnetospirillum magneticum* (ATCC 700264), Wolfe’s Vitamin Solution and Wolfe’s Mineral Solution were purchased from American Type Culture Collection (Manassas, VA). 1,2-dipalmitoyl-sn-glycero-3-phosphocholine (DPPC) and 1,2-distearoyl-sn-glycero-3-phosphoethanolamine-N-[carboxy(polyethylene glycol)-2000] (sodium salt) (DSPE-PEG2000-COOH) were purchased from Avanti Polar Lipids, Inc. (Alabaster, AL). Matrigel (HC) and penicillin/Streptomycin (P/S) were purchased from Corning (Corning, NY). Fetal bovine serum (FBS) was purchased from BioWest (Nuaillé, France), 1-ethyl-3-[3-(dimethylamino)propyl]-carbodiimide (EDC) was purchased from TCI EUROPE (Eschborn, Germany), and Hoechst 33342 was purchased from Invitrogen (Burlington, CA). Calcein and 4% formaldehyde (HistoFix 4%) were purchased from Carl Roth (Karlsruhe, Germany). VE-cadherin rabbit monoclonal primary antibody and Alexa488-conjugated anti-rabbit IgG secondary antibody were obtained from Cell Signalling Technology (Danvers, MA). Potassium phosphate, succinic acid, tartaric acid, sodium nitrate, ascorbic acid, sodium acetate, resazurin sodium salt, gelatin, epidermal growth factor (EGF), hydrocortisone, Hanks’ Balanced Salt solution (HBSS), dimethyl sulfoxide (DMSO), Lucifer yellow CH dipotassium salt, sepharose CL-2B, bovine serum albumin (BSA), 3,3′-Dioctadecyloxacarbocyanine perchlorate (DiO), phosphate-buffered saline (PBS), Triton X-100 and N-hydroxysulfosuccinimide (sulfo-NHS) were all acquired from Sigma-Aldrich (St. Louis, MO). Dulbecco’s Modified Eagle’s Medium (DMEM), McCoy’s 5A medium, L-glutamine, CellTrace Far-Red cell proliferation kit, agarose and MCDB131 medium were acquired from Thermo Fisher Scientific (Waltham, MA).

### Mammalian cell culture

Caco-2 cells were cultured in high glucose DMEM supplemented with 20% FBS and 1% P/S. HMEC-1 cells were cultured in MCDB131 supplemented with 10% FBS, 1% P/S, 10 mM L-glutamine, 1 µg/mL hydrocortisone, 10 ng/mL EGF. MCF-7 cells were cultured in high glucose DMEM supplemented with 10% FBS and 1% P/S. HCT 116 cells were cultured in McCoy’s 5A medium supplemented with 10% FBS and 1% P/S. All cell lines were incubated at 37 °C with 5% CO_2_ in a humidified atmosphere. Cells were harvested at 80% confluency and cell density was determined using a hemocytometer. The required number of cells was seeded for each experiment as specified. One day before experiments, media was replaced with antibiotic-free media.

### Bacteria culture

Cultures of *M. magneticum* strain AMB-1 were grown in revised magnetic spirillum growth medium (MSGM, ATCC Medium 1653) which contained the following per liter of distilled water: 5.0 mL of Wolfe’s Mineral Solution, 0.68 g of potassium phosphate, 0.37 g of succinic acid, 0.37 g of tartaric acid, 0.12 g of sodium nitrate, 0.035 g of ascorbic acid and 0.05 g of sodium acetate. The final pH was adjusted to 6.75 with 1 M NaOH before autoclaving. Prior to use, Wolfe’s Vitamin Solution (1000x) and 10 mM ferric quinate (200x) were added to the culture media. Incubation occurred anaerobically at 30 °C and cultures were passaged every 5 to 7 days.

### Caco-2 monolayer culture and permeability assay

Caco-2 cells were seeded at a density of 1 × 10^5^ cells/cm^2^ on 12-well Transwell inserts (3.0 µm pore size, Corning). Cells were cultured on the inserts for 16-21 days and media was exchanged every second day. A Lucifer yellow paracellular permeability assay was performed to determine the integrity of the established Caco-2 monolayer. Prior to and after each assay, cell monolayers were washed three times with HBSS. The basolateral compartment of a well plate was filled with 1.5 mL of 1% DMSO in HBSS and 0.5 mL of 1 mg/mL Lucifer yellow in 1% DMSO/HBSS was added to the apical compartment. After one hour incubation at 37 °C, the fluorescence intensity (FI) of the solutions from the apical and basolateral compartments were measured at 485/535 nm using a Spark multimode microplate reader (Tecan). The percent rejection of Lucifer yellow was calculated as follows:

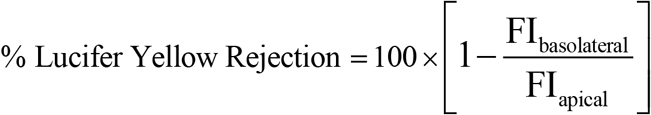

Transwell inserts with a Lucifer yellow rejection value greater than 99% were used for further experiments. After each experiment, the Lucifer yellow assay was repeated to confirm that the integrity of the monolayer was maintained.

### Integrated imaging and wireless magnetic control platform

A small-scale arbitrary magnetic field generator consisting of eight electromagnets arranged in a single hemisphere was used to apply rotating magnetic fields (Magnebotix, Zurich, Switzerland). The system was integrated with an inverted spinning disk confocal microscope (Nikon Eclipse Ti2 with Yokogawa CSU-W1 unit and Hamamatsu C13440-20CU Digital CMOS camera), which was used for all imaging. Samples were positioned between the objective lens and the hemisphere of the electromagnets.

### MTB translocation across Caco-2 cell monolayers

Transwell inserts with Caco-2 monolayers were placed into PDMS wells (Ø = 22 mm) containing 1.5 mL media. MTB at a concentration of 1 × 10^8^ MTB/cm^2^ was added to the apical chamber and samples were exposed to either a directional magnetic field of 12 mT or a localized rotating magnetic field of 12 mT with frequencies between 1 and 20 Hz. A magnetic force generated by the superposition of moderate gradients of 1 T/m was applied along –z. After one hour, the bacteria concentration in the basolateral chamber was determined by live counting in a hemocytometer. Bacteria concentrations for actuated samples were normalized to unactuated controls.

### Inductive detection of MTB

Inductive detection of MTB was performed using a custom-built RMF magnetometer. Two pairs of nested Helmholtz coils were wound using enamel-coated copper wire (Ø = 0.8 mm) and housed in a 3D printed frame (fig. S4). The inner set of coils contained 256 turns of wire and the outer coils consisted of 265 turns. Phase shifted sinusoidal signals were generated (Digilent Analog Discovery 2) and amplified (Crown D-150A Series II), resulting in a sinusoidal waveforms in each set of Helmholtz coils to produce a circularly rotating magnetic field. Inductive field probes for each pair of coils were placed in the central workspace along with sense and compensation coils. Signals from the sense and compensation coils underwent differential amplification and the residual signal from the magnetized sample was acquired with an oscilloscope (Keysight DSOX2004A). All samples contained 100 μL of MTB at approximately OD_600_ = 1.5, suspended in MSGM. Measurements were collected at field amplitudes of 12 mT and 20 mT at a frequency range between 12 Hz and 80 Hz. A detailed description of the setup and the associated data processing pipeline can be found in the supplementary materials.

### Computational model of endothelial translocation

A numerical simulation in COMSOL Multiphysics was implemented to study the translocation of microstructures across an endothelial barrier under different conditions. MTB were modeled as rigid ellipsoids possessing a magnetic dipole moment using dimensions obtained from Multisizer data collected after *in vitro* translocation experiments. Endothelial cells were modeled as hyperelastic materials and stochastic opening of cell-cell contacts was incorporated to model dynamic gap formation that occurs in endothelial cell monolayers. The effect of the surrounding fluid was modelled through linear and rotational drag coefficients assuming Stokes flow regime. A time-varying external magnetic field was applied, and this was balanced with forces and torques arising from hydrodynamic and mechanical interactions with the viscous fluid and adjacent substrate. Further details can be found in the Supporting Information.

### Liposome preparation and characterization

Fluorescently-labeled carboxylated liposomes were prepared from a total of 14 μmol of lipids using thin-film hydration. For imaging, DPPC, DSPE-PEG2000-COOH (5 mol%) and DiO (0.5 mol%) were combined in 0.5 mL chloroform and dried to a thin film under nitrogen with further vacuum desiccation overnight at room temperature. The lipid thin film was hydrated with 1 mL PBS and placed in a water bath for 1 h at 50 °C with continuous stirring. Liposomes were downsized using sequential extrusion on a heating block (Avanti) at 50 °C. Liposomes were passed 21-times each through a 400 nm followed by a 200 nm polycarbonate membrane (Whatman). For flow cytometry, DPPC and DSPE-PEG2000-COOH (5 mol%) were combined to form a thin film as previously described. The film was rehydrated with 1 mL calcein solution (35.2 mM) and downsized as described. Unencapsulated calcein was removed by size exclusion chromatography using a Sepharose CL-2B column. The average diameter, size distribution and zeta potential of the liposomes was determined by dynamic light scattering (DLS) (Litesizer 500, Anton Paar).

### MTB staining and covalent coupling of liposomes

MTB cells at a concentration of 1.5 × 10^8^ cells/mL in PBS were stained using the CellTrace Far-Red cell proliferation kit. A stock solution was prepared according to the manufacturers protocol and 2 µL was added to the MTB suspension. Cells were incubated at room temperature for 20 min, protected from light with gentle agitation. After incubation, 10 µL of 1% BSA was added to the cell suspension for 5 min to remove free dye. Cells were pelleted and resuspended in PBS.

Bioconjugation was then achieved through carbodiimide coupling using an adapted protocol (*20*). Briefly, activation of carboxylated liposomes was accomplished by incubating 300 μL of liposome solution with EDC and sulfo-NHS (EDC/NHS/DSPE-PEG-COOH = 30:30:3, mol/mol) for 20 min in PBS (pH 5.5) at room temperature with gentle agitation. The activated liposomes were subsequently incubated with 1.5 × 10^8^ MTB cells for 2 hours at room temperature with gentle agitation. Unbound liposomes were separated from MTB-liposome conjugates (MTB-LP) using a two-dimensional magnetic field and the sample was redispersed in PBS (pH 7.4).

### Quantification of liposome binding by flow cytometry

For quantitative analysis of liposome binding, MTB-LP conjugates were analyzed using flow cytometry (BD LSRFortessa) with a 488 nm excitation laser and 530/30 filter. Untreated MTB cells and unbound liposomes were used as controls and the fluorescence emission of 10,000 events was recorded. Data was analyzed using FlowJo (Tree Star) and appropriate gates and controls were used to generate density plots and histograms of each sample.

### HMEC-1 monolayer culture and immunofluorescence staining

HMEC-1 cells were seeded at a density of approximately 1 × 10^5^ cells/cm^2^ on 12-well Transwell inserts (3.0 µm pore size) and cultured on the inserts for 2 days. HMEC-1 monolayers were fixed in 4% paraformaldehyde for 15 min and permeabilized in 0.1% Triton X-100 for 10 min at room temperature. Fixed cells were blocked with 1% BSA in PBS for 1 hour, followed by overnight incubation at 4 °C with a VE-cadherin rabbit monoclonal primary antibody (diluted 1:500 in blocking buffer). The monolayers were then incubated for 1 hour at room temperature in an Alexa488-conjugated anti-rabbit IgG secondary antibody (diluted 1:1000 in blocking buffer). Staining for DNA was performed with Hoechst 33342. Monolayers were incubated in Hoechst 33342 at a final concentration of 10 µg/mL in PBS for 10 min at room temperature. The membranes were then detached from the Transwell inserts, mounted on glass coverslips and imaged.

### Translocation across HMEC-1 cell monolayers

HMEC-1 monolayers cultured on Transwell inserts were placed into PDMS wells (Ø = 22 mm) containing 1.5 mL media. MTB-LP at a concentration of 1 × 10^8^ MTB-LP/cm^2^ was added to the apical chamber and exposed to either a directional magnetic field of 12 mT or a localized rotating magnetic field of 20 mT and 24 Hz. After one hour, the concentration and size distribution of MTB-LP in the basolateral chamber was determined using a multisizer (4e Coulter Counter, Beckman Coulter). For comparison, liposomes were added to monolayers at a concentration corresponding to the liposomes on the conjugates. After 1 hour, the solutions from the apical and basolateral compartments were collected and measured at 485/535 nm using a Spark multimode microplate reader (Tecan).

Prior to and after each assay, LY rejection (as previously described) and transendothelial electrical resistance (TEER) measurements for performed using a EVOM3 Volt/Ohmmeter (World Precision Instruments). TEER was calculated using following the equation:

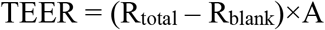

where R_total_ is the measured resistance from a Transwell insert with a cultured monolayer, R_blank_ is the the resistance of a Transwell insert without a monolayer, and A is the area of Transwell insert (1.12 cm^2^).

### MCF-7 spheroid formation and staining

MCF-7 tumor spheroids were formed and cultured in 96 well ultralow adhesion plates (Corning). Cells were seeded at a density of 10,000 cells/well in 100 µL of media. The well plates were centrifuged at 1000 x g for 10 min and incubated in a humidified atmosphere with 5% CO_2_ at 37 °C. Tumor spheroids grew to approximately 500 µm in diameter after 3 days. Prior to experiments, staining for DNA was performed with Hoechst 33342. Spheroids were incubated for one hour at 37°C with Hoechst 33342 at a final concentration of 5 µg/mL in media.

### MTB-LP accumulation in MCF-7 spheroids

MCF-7 spheroids were transferred to PDMS wells (Ø = 6 mm) containing 5 × 10^7^ MTB-LP in 100 µL of media. Spheroids were subsequently imaged under magnetic actuation at 20 mT and 24 Hz. After one hour, spheroids were washed thoroughly with PBS and incubated in ultralow adhesion well plates for up to 6 days at 37 °C with 5% CO_2_ in a humidified atmosphere.

### Quantification of MTB-LP accumulation in MCF-7 spheroids

Analysis and processing of captured image stacks was performed in ImageJ (NIH). For each spheroid, a summation of consecutive Z-plane images in the first 100 μm was performed for each time point (fig. S9A). Regions of interest (ROIs) were defined using the Hoechst image stack by tracing a contour around the spheroid at 0 and 100 µm. Intermediate ROIs were defined using interpolation, with approximately 3 µm radial increments (fig. S9B). These ROIs were then applied to the corresponding DiO liposome image stack and fluorescence intensity measurements were performed. Fluorescence values were normalized to the fluorescence of the surrounding media (fig. S9C).

To generate fluorescence intensity distributions at a depth of 80 μm, rectangular regions of interest were defined at the boundaries of the spheroid. The average pixel intensity values along the x-axis were normalized to the minimum and maximum intensity across all the spheroids to allow comparison, using the following equation:

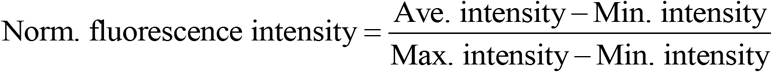

Normalized fluorescence intensity was plotted against normalized x-axis distance to obtain the MTB-LP distribution in the spheroids.

### *In vivo* magnetic actuation experiments

Female BALB/c nude mice (6-8 weeks, Charles River) were acclimatized for 3 weeks and inoculated subcutaneously in the hind flank with 5×10^6^ MCF-7 cells suspended in 8 mg/mL Matrigel at a volume of 100 μl. When tumor diameters exceeded 5 mm, mice were blindly randomized into various groups with a minimum of 3 mice per group. Tail-vein (intravenous) injections of MTB stained with a far-red proliferative dye were administered at a concentration of 1×10^9^ in 100 μl PBS. Mice were placed under anesthesia in the absence of magnetic actuation (control) or with tumors positioned in the workspace of the magnetic field generator and exposed to actuation at 20 mT and 14 Hz for 1 h (RMF). Mice were then returned to the cage for 24 h, after which the mice were euthanized and the tumor and major organs were harvested. For semiquantitative biodistribution analysis, far-red fluorescence signals were measured using an *ex vivo* fluorescence imaging system (Sapphire Biomolecular Imager, Azure Biosystems).

Harvested tumors were weighed and homogenized using a gentleMACS tissue dissociator (Miltenyi Biotec) (C-tubes) before being cultured for 8 days under the MTB culture conditions described previously. C_mag_ measurements were performed to quantify the magnetic responsiveness of the samples. Optical density (OD) at 600 nm was measured with a magnet placed parallel (OD_||_) and then perpendicular (OD_⊥_) to the light path. The C_mag_ value was calculated as the fraction of OD_||_ and OD_⊥_.

### Histology

Spheroids and harvested tumors were fixed 144 h in 4% formaldehyde for 1 hour and washed three times with PBS. For spheroids, a 5% (w/v) agarose solution in DI water was mixed with a 5% (w/v) solution of gelatin at a 1:1 ratio, and the fixed spheroids were embedded in the resulting hydrogel. After dehydration in graded alcohol and isoparaffin (LOGOS J, Milestone Medical), spheroid-containing hydrogels or tumors were embedded in paraffin and sections of 5 – 10 μm were cut using a microtome (HM 355S, Microm AG). The sections were stained with Hoechst 33342 at a concentration of 5 µg/mL in DI water and the slides were mounted and imaged.

### Statistical Analysis

All values are expressed as the mean ± standard deviation (SD) of at least three replicates, unless otherwise stated. Statistical analysis between two groups was performed using an unpaired Student’s t-test and for comparisons of more than two groups, analysis of variance (ANOVA) with Tukey’s post hoc analysis was performed. A p-value < 0.05 was considered statistically significant. Plotting of graphs and statistical analysis were performed using GraphPad Prism 8.0 software.

## Supplementary Materials

Supplementary Text

Fig. S1. Comparison of MTB translocation across Caco-2 monolayers at varying out-of-plane rotational frequencies.

Fig. S2. Lucifer yellow rejection values for Caco-2 monolayers.

Fig. S3. RMF magnetometer for inductive detection of MTB.

Fig. S4. MTB size distributions before and after actuation.

Fig. S5. Size distribution of liposomes.

Fig. S6: Bioconjugation of carboxylated liposomes to MTB.

Fig. S7. Assessment of MTB colonization in HCT 116 tumor spheroids.

Fig. S8. Plots of expected field magnitude and gradients

Fig. S9: Quantification of fluorescence intensity in spheroids.

Table S1: Physicochemical characterization of liposomes.

Movie S1: Modeling of liposome diffusion across endothelial monolayer.

Movie S2: Computational modeling of MTB-LP transport across modelled endothelial monolayer using DMF.

Movie S3: Computational modeling of MTB-LP transport across modelled endothelial monolayer using RMF.

Movie S4: Actuation of MTB on HMEC-1 monolayer

Movie S5: MTB actuation into MCF-7 spheroid

## Acknowledgments

The Flow Cytometry Core Facility at ETH Zurich provided technical support for flow cytometry measurements. The authors thank Dr. David C. Bono for useful input for the magnetometer design, Lucas Amoudruz for helpful discussions on computational modeling of MTB, and Lucien Stöcklin for his assistance with the inductive detection experiments. The authors also thank Dragana Ristanovic for helpful discussions and assistance with editing the manuscript. TG was supported by a Swiss Government Excellence Scholarship.

## Author contributions

Conceptualization: SS, TG, NM, MGC, VL

Methodology: SS, TG, NM, MGC

Investigation: TG, NM, MGC, TT

Visualization: TG

Supervision: SS

Writing – original draft: TG

Writing – review & editing: SS, TG, MGC, NM, TT, VL

## Data and materials availability

All data needed to evaluate the conclusions in the paper are present in the paper and/or the Supplementary Materials. Additional data related to this paper may be requested from the authors.

## SUPPLEMENTARY MATERIALS

### MTB propulsive force and torque-based force calculations

MTB motion is dependent on the propulsive force (*F*_*P*_) generated by their flagella and the fluidic drag force (*F*_*D*_). At constant velocity, this relationship is governed by:

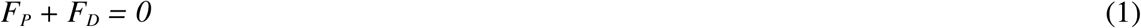

When traveling at constant velocity at low Reynolds numbers the propulsive force is equal and opposite to the drag force which is defined as:

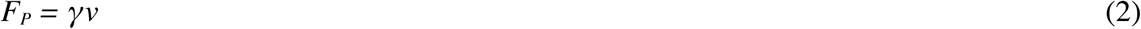

where *v* is the linear velocity and γ is the linear drag coefficient given by (*55*):

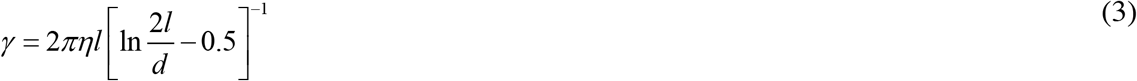

Thus, for a velocity range of 19 - 49 µm/s the propulsive force is estimated to fall between 0.14 and 0.35 pN when the dynamic viscosity of the medium (*η*) is approximately 1 × 10^−3^ Pa·s, and the length and diameter of the cell are *l* = 1.8 µm and *d* = 0.45 µm, respectively (fig. S1).

In contrast, when MTB are exposed to an externally rotating magnetic field, they experience a magnetic torque, *τ*_*M*_ *=* |***m × B***| *= m B sin θ*, dependent on the magnetic moment (*m*) of the MTB and the magnetic field strength (*B*). The opposing hydrodynamic torque τ_H_ defines the steady state lag angle (*θ*) between the MTB and magnetic field and is governed by *τ*_*H*_ = *a ω*, where the angular frequency (*ω*) is equal to the frequency of magnetic actuation under synchronous rotation. The rotational drag coefficient (*α*) is obtained by (*56*):

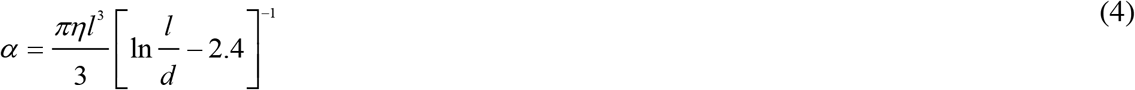

Thus, the force (*F*_*T*_) generated by this torque-based motion can be determined using the following equation:

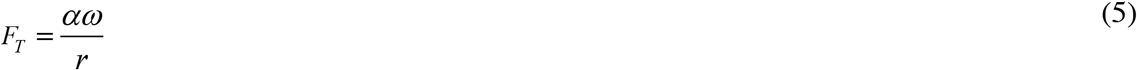

For angular frequencies between 10 – 25 Hz and distance from the axis of rotation (*r*) of 0.9 µm, the force generated ranges from 0.8 – 1.8 pN.

### RMF magnetometer for inductive detection of MTB

Inductive detection of MTB was performed using a custom-built RMF magnetometer. Two pairs of nested Helmholtz coils housed in a 3D printed frame were used to produce a circularly rotating magnetic field (fig. S4). Field magnitude was determined by measuring current with a shunt resistor and calibrating under constant field to a Hall probe (Metrolab THM1176), which agreed well with the expected geometric factor. Field measurements were ultimately performed with orthogonal inductive field probes (100 turns) placed in the central workspace along with sense and compensation coils (each 500 turns). Signals from the sense and compensation coils underwent differential amplification (Analog Devices EVAL-ADA4625-1) and the residual signal from the sample was acquired by an oscilloscope (Keysight DSOX2004A) under signal averaging (N = 32).

Measurements were collected at field amplitudes of 12 mT and 20 mT over a logarithmically spaced frequency range between 12 Hz and 80 Hz. Three measurements per frequency and amplitude were collected for blank samples containing MSGM and samples containing MTB. An inductive background signal at the same frequency and at 2 mT was collected both before and after every sample measurement. This approach assumes that the response of the bacteria at 2 mT and the investigated frequencies will be far smaller than their response at higher amplitudes, an assumption motivated by previous studies (*37*). Prior to conducting measurements at each condition, inductive background was minimized by minutely mechanically adjusting the angle between the sense and compensation coil to balance phase and using the potentiometer to minimize the output signal of the amplifier. After a sample or blank was loaded, care was taken not to touch the device or otherwise disturb background cancellation.

During subsequent data processing, any offset arising from amplification was subtracted and only the time-varying components of the acquired signals were considered. A linear combination of the background signals of the two field probes was fitted to the 2 mT magnetometer signals. This uses the orthogonal signals from the field probes as references for reconstructing residual inductive signal. Estimates of the background based on the measurements at 2 mT conducted immediately prior to and after measurement at the target amplitudes were then subtracted from the blank and MTB samples, providing initial and final bounds on the sample and blank signals. Each of the bounded blank signals (six) were then subtracted from each of the bounded MTB signals (six) to form a set of 36 signals. Because background subtraction is likely dominate error in this case, this set of 36 signals can be taken to represent the variation of equally theoretically valid background-subtracted signals.

From this collection of 36 bounding signals, the mean signal generated by the MTB was found and bootstrapping was performed to provide 95% confidence intervals to quantify the uncertainty in the data presented in Figure 2A. Using a cosine function fitted to the in-phase field probe, integration was used to separate each of these 36 signals into quantities proportional to the in-phase and out-of-phase components of the magnetization. Dividing by frequency and field magnitude yielded quantities proportional to susceptibility that can be compared to each other over the range of frequencies and amplitudes. Bootstrapping was also used on these sets of values to provide a 95% confidence interval on the resulting values for real and imaginary susceptibility depicted in Figure 2B.

Predictions of the susceptibility values for MTB where made by plotting the normalized real (χ’) and imaginary (χ’’) components of susceptibility at 12 and 20 mT using the following equations:

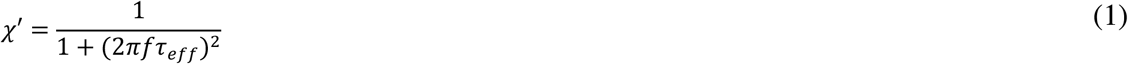

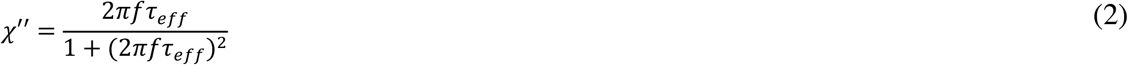

where f is the frequency of rotation and τ_eff_ is the effective Brownian relaxation time given by:

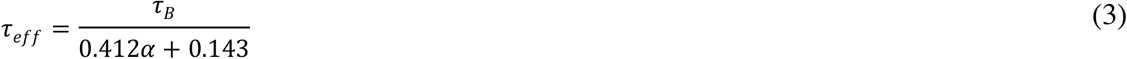

The Langevin parameter, α, is defined by:

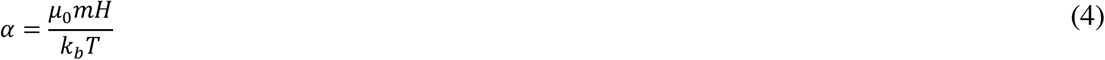

where, *μ*_*0*_ denotes vacuum permeability, *m* is the magnetic moment, *H* is the external field magnitude, *k*_*b*_ is Boltzmann’s constant and the temperature is T = 298 K.

The characteristic zero-field Brownian relaxation, *τ*_*B*_ time is given by:

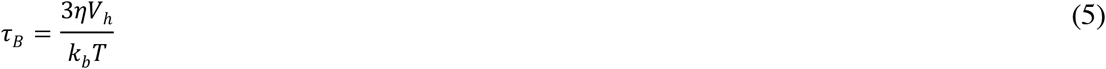

where *η* is the viscosity of the fluid and *V*_*h*_ is the hydrodynamic volume.

### Computational modeling of MTB-LP transport across endothelial monolayers

To study the transmigration mechanism of MTB across HMEC monolayers, Finite Element Method (FEM) was employed to solve the governing equations using COMSOL Multiphysics. The cell monolayer was modeled in 2D featuring a few adjacent endothelial cells forming a sealed barrier between upper and lower compartments. Endothelial cells were modeled as hyperelastic material with shear modulus of 1 kPa (*33, 34*). The dimensions of the cells in the monolayer was adopted from Arefi et al. (*34*). Considering the relative stiffness of gram-negative bacteria compared to endothelial cells, MTB were treated as rigid bodies. MTB were modeled as ellipsoids initially present in the upper compartment possessing a rigid dipole moment along their long axes. TEM images along with multisizer data (fig. S1) were used to estimate the average size of a single bacterium. Assuming 0.45 um for the short axis and using the diameter of equivalent volume from multisizer measurements, one could estimate the long axis of MTB to be approximately 1.8 um.

Motion of micron-sized objects in a fluid close to a wall is well studied as surface walkers under low Reynolds number flows (*57*–*59*). The free body diagram of single bacterium illustrates the balance of acting forces and torques in such an environment:

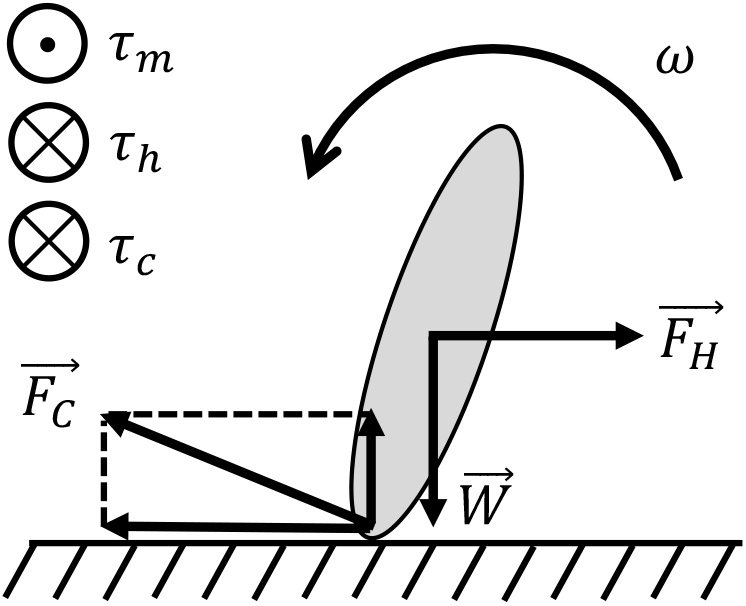

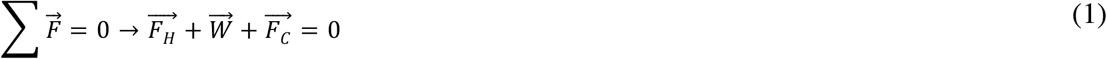

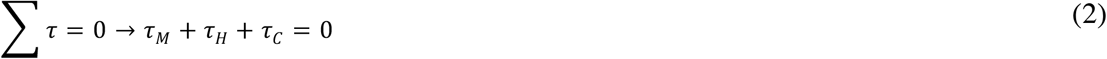

*F*_*H*_ indicates the viscous drag, *W* is the gravity force, and F_C_ is the reaction force from the substrate including both normal and frictional components. A mass density of *ρ* = 200 *kg / m*^*3*^ was assumed for the bacteria to take the effect of buoyant force into account (*60*). In the conservation of angular momentum equation, *τ*_*M*_ represents magnetic torque, *τ*_*H*_ is hydrodynamic resistant torque, and *τ*_*C*_ indicates the torque coming from the interaction with the substrate and is given by:

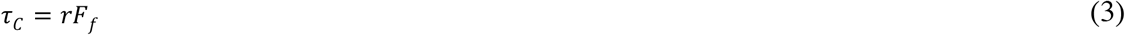

where the distance from the axis of rotation *r* = 0.9 µm and *F*_*f*_ is the friction force.

Magnetic torque arising from the lag between the magnetic dipole moment of MTB and external field is given by:

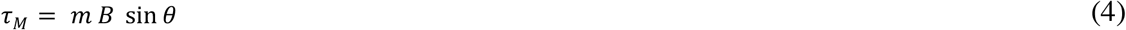

where *m* is the magnetic dipole moment of MTB which is estimated from VSM measurements (*37*), *B* represents the external magnetic field, i.e. 20 mT, and *θ* is the phase lag between two vectors.

Stokes flow theorem correlates hydrodynamic interactions in terms of linear and rotational viscous drag with corresponding linear and angular velocities:

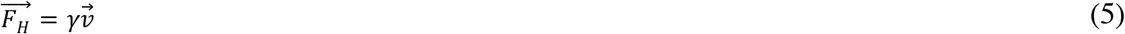

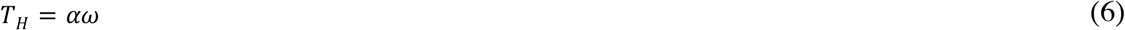

where *γ* and *α* are the linear drag coefficient rotational drag coefficient, respectively. Analytical expressions and empirical data were used to estimate the values of these these parameters which were set to *γ* = 7.1 × 10^−9^ *N*. *s/m* and *α* = 7.6 × 10^−20^ *N. m. s/rad* in the simulations.

The velocity dependence of friction between two microscale objects has previously been reported (*61*). This feature was incorporated into the friction model in which the slip and no slip regime was determined by a critical friction force depending on normal traction and velocity difference between two objects at the point of contact. Numerically, the penalty method was employed to resolve the mechanical contact which is based on insertion of a spring between two objects, only active in the case of compression.

Rigid Body node of the Solid Mechanics interface was used to solve the above governing equations. Hydrodynamic interactions were then modeled as linear and rotational viscous damping acting on the rigid body.

In order to model the dynamic gaps forming stochastically between adjacent cells, a set of random parameters were generated for each simulation. These values determined the opening time of the junction and size of the gaps. Random gap size was within the range of 1.5 to 2.5 um (*35*) and the gap lifetime was set to 160 s (*36*). The overall simulation time was selected in a way that encompasses opening incidences of all gaps. Considering the size of the computational domain and translational motion of the bacteria, the simulation was carried out for 500 s.

### Liposome characterization and bioconjugation of carboxylated liposomes to MTB

Bioconjugation of fluorescently-labelled liposomes to the MTB cell surface was achieved through carbodiimide mediated amidation. This coupling reaction relies on the terminal amine groups present in phospholipids, lipopolysaccharides, and various surface protein assemblies on the MTB cell membrane (*62*). In comparison to affinity-based conjugation, covalent crosslinking is a robust technique which produces a stable chemical bond for *in vitro* and *in vivo* applications (*63, 64*).

Carboxylated liposomes fabricated using thin film hydration were found to have an average size of 184.2 ± 3.2 nm and a polydispersity index of 0.142 ± 0.055, indicating that the liposomes were monodisperse (fig. S5, Table S1). Zeta potential analysis showed that the liposomes had a negative net surface charge of 39.8 ± 0.9 mV, confirming successful incorporation of charged carboxylic groups on the outer membrane of the liposomes.

The reactive carboxy groups on the liposomes were then covalently coupled to the primary amines on the MTB membrane using NHS chemistry. Flow cytometry was performed to confirm successful conjugation. Univariate histograms show a clear distinction between bacteria with low relative fluorescence and MTB-LP, which extend to higher relative fluorescence intensity values (fig. S6). The mean fluorescence intensity of the conjugates was approximately 380-fold higher than that of bacteria cells, signifying that liposomes were conjugated to the bacteria.

## Supplementary Figures

**Fig. S1.**
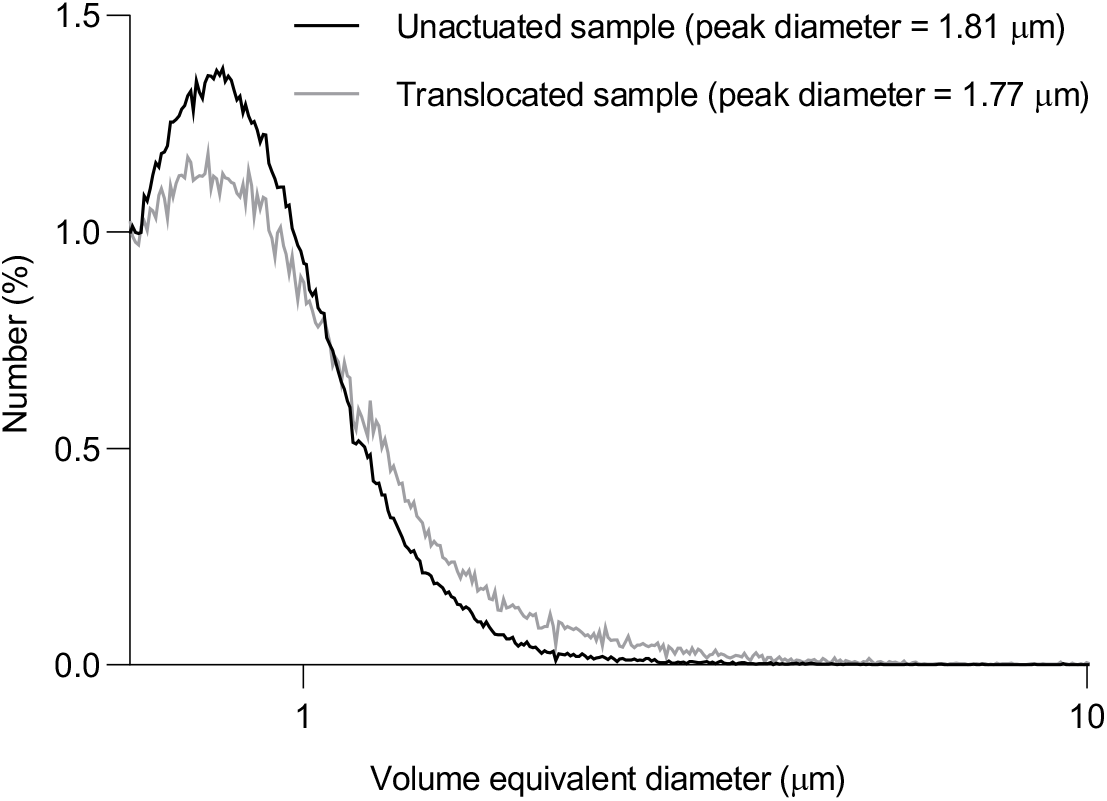
MTB size distributions before and after actuation. Average size distribution and average peak diameter values of MTB obtained by a Coulter counter. The plotted values are for the diameter of a sphere with equivalent volume and were used to estimate dimensions of the bacterial body (*n* = 3; mean).

**Fig. S2.**
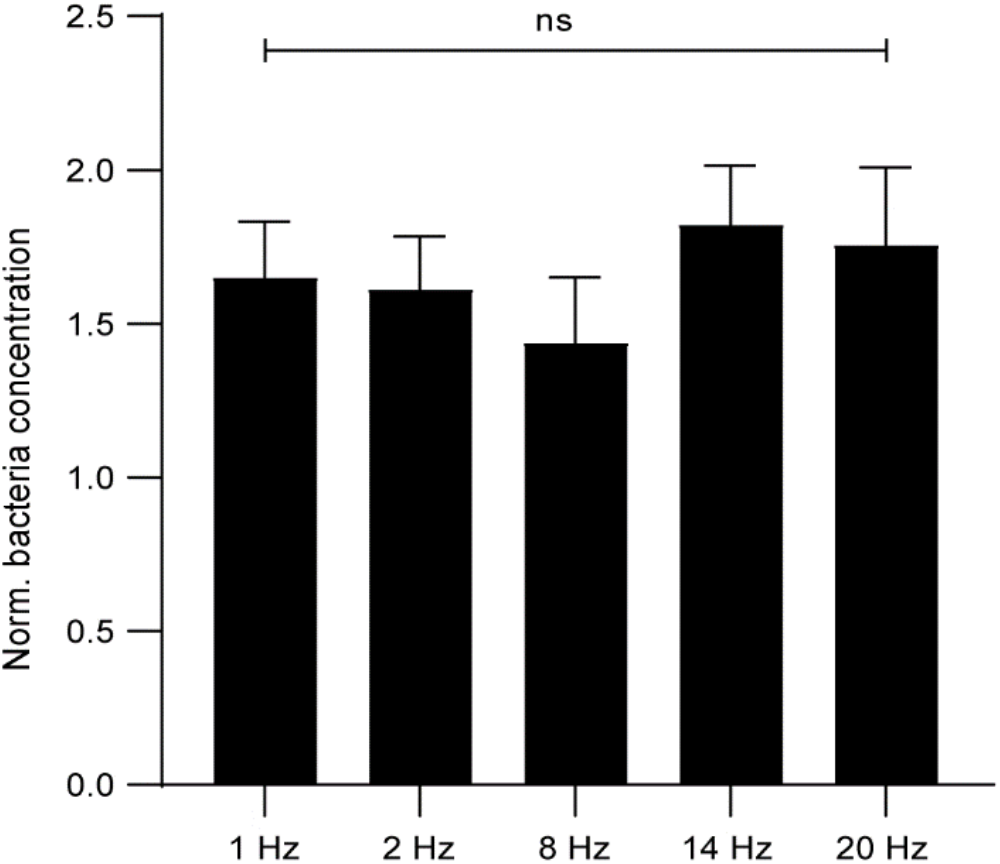
Comparison of MTB translocation across Caco-2 monolayers at varying out-of-plane rotational frequencies. Bacteria concentration was normalized to respective unactuated controls. No significance between all conditions ((*n* = 3; mean ± SD; ANOVA).

**Fig. S3.**
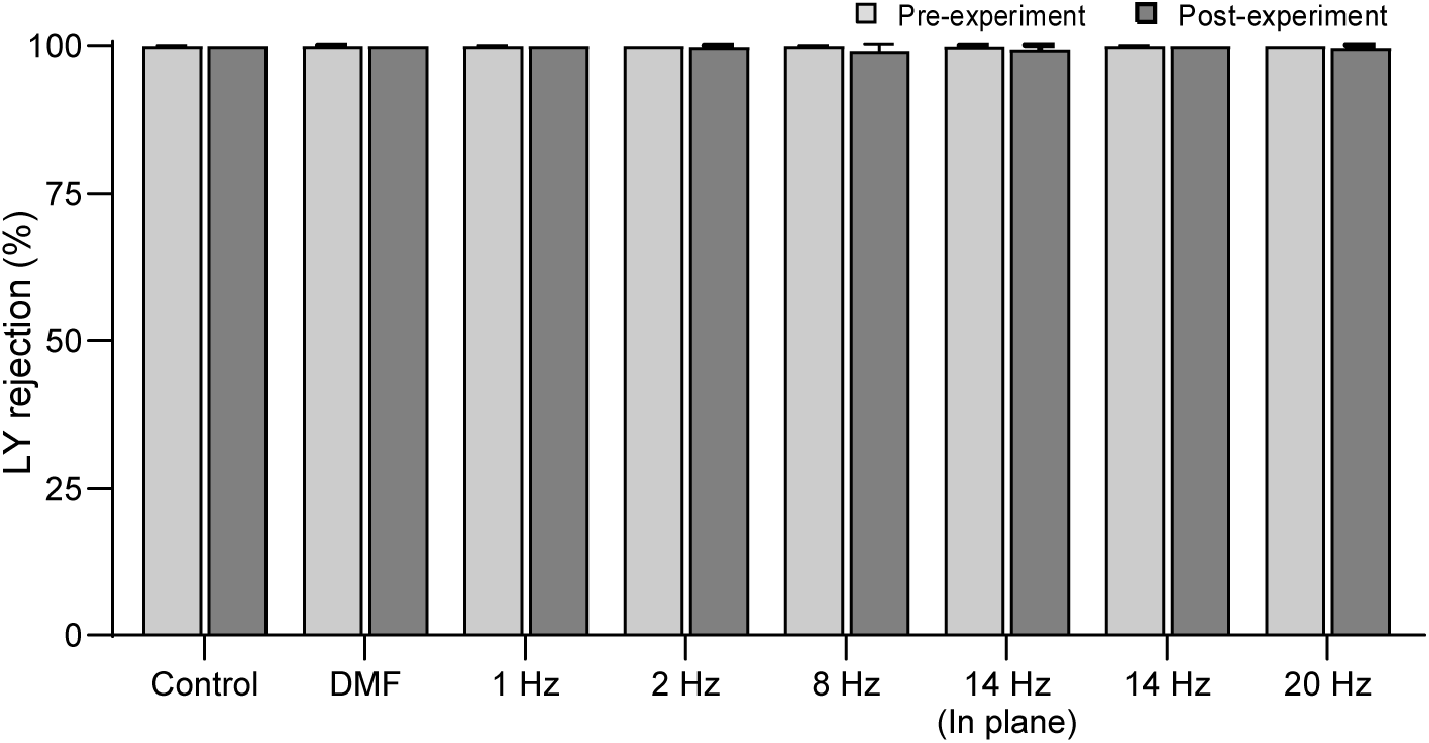
Lucifer yellow rejection values for Caco-2 monolayers. Pre- and post-experimental Lucifer yellow (LY) rejection values for all tested conditions. Experiments were performed at a magnetic field strength of 12 mT for all conditions (*n* = 3; mean ± SD).

**Fig. S4.**
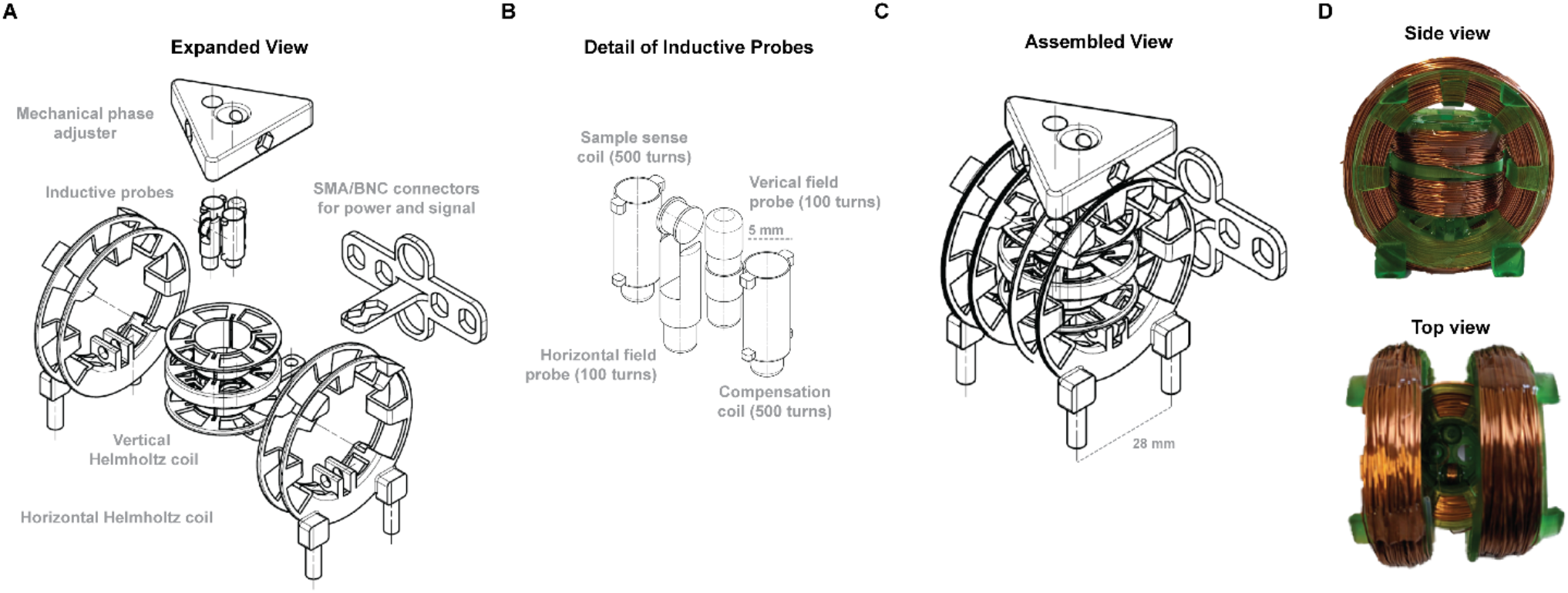
RMF magnetometer for inductive detection of MTB. (**A**) Technical drawing of RMF magnetometer frame and components. (**B**) Detailed drawing of inductive probes. (**C**) Technical drawing of assembled RMF magnetometer. (**D**) Images of side view (top) and top view (bottom) of assembled RMF magnetometer.

**Fig. S5.**
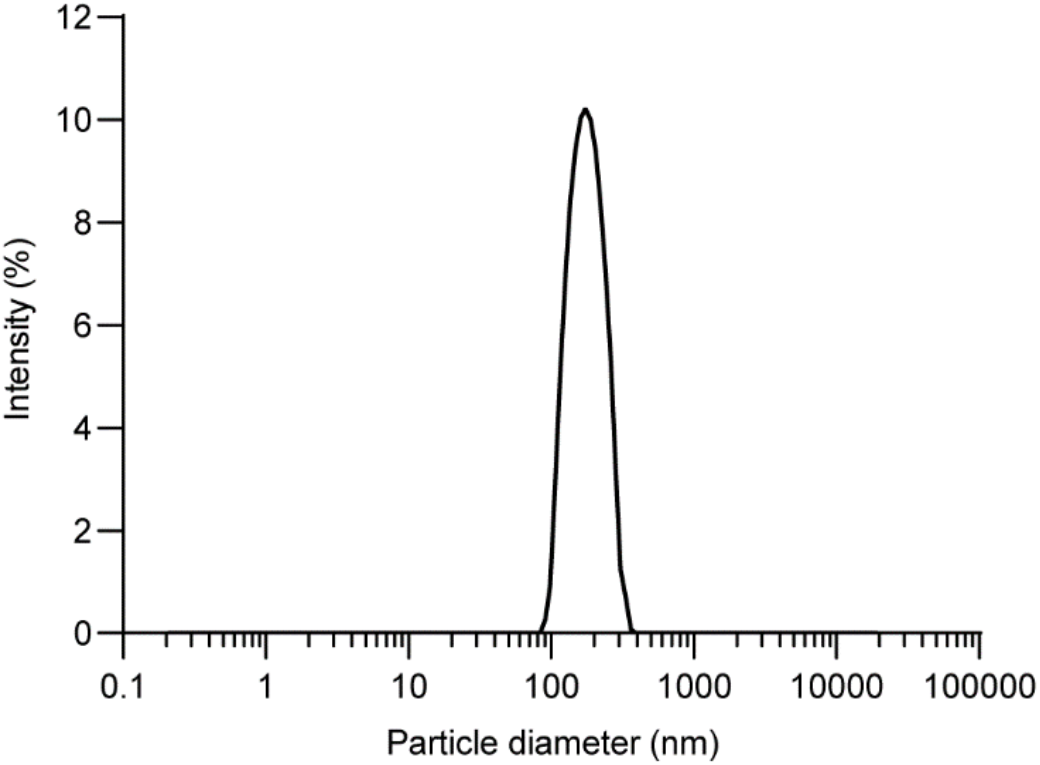
Size distribution of liposomes. Data collected using dynamic light scattering (DLS) (*n* = 3; mean).

**Fig. S6:**
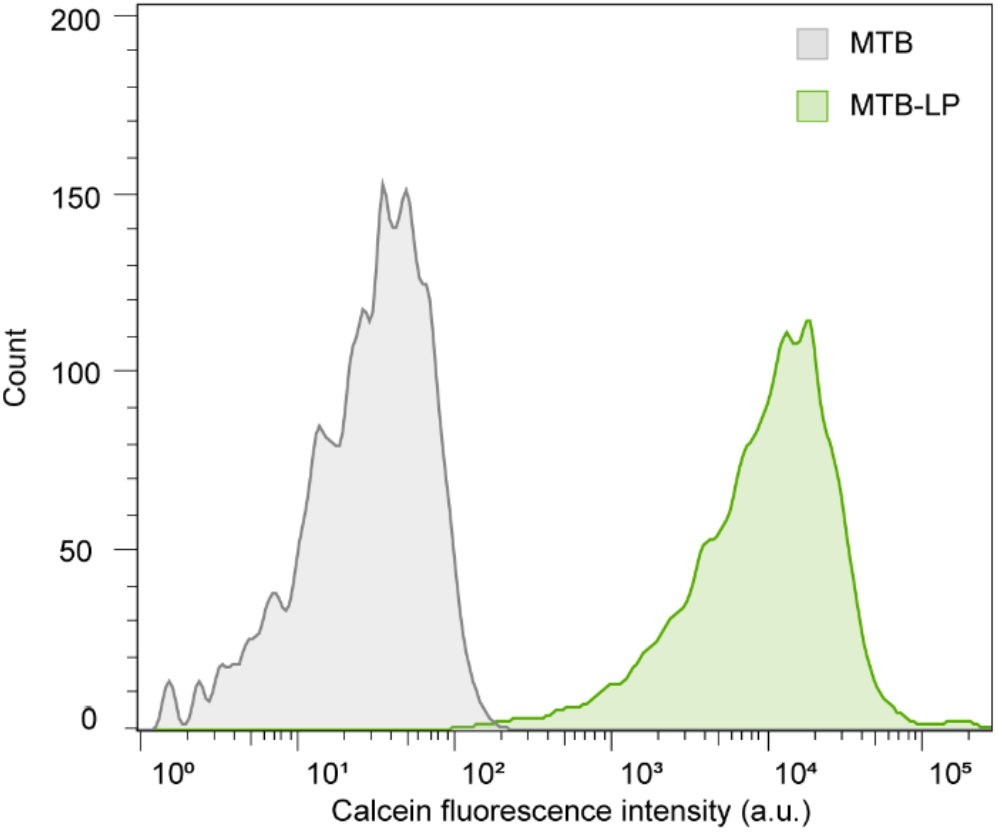
Bioconjugation of carboxylated liposomes to MTB. Representative flow cytometry histograms of AMB-1 bacteria (MTB), and conjugates of MTB and calcein-loaded liposomes (MTB– LP).

**Fig. S7.**
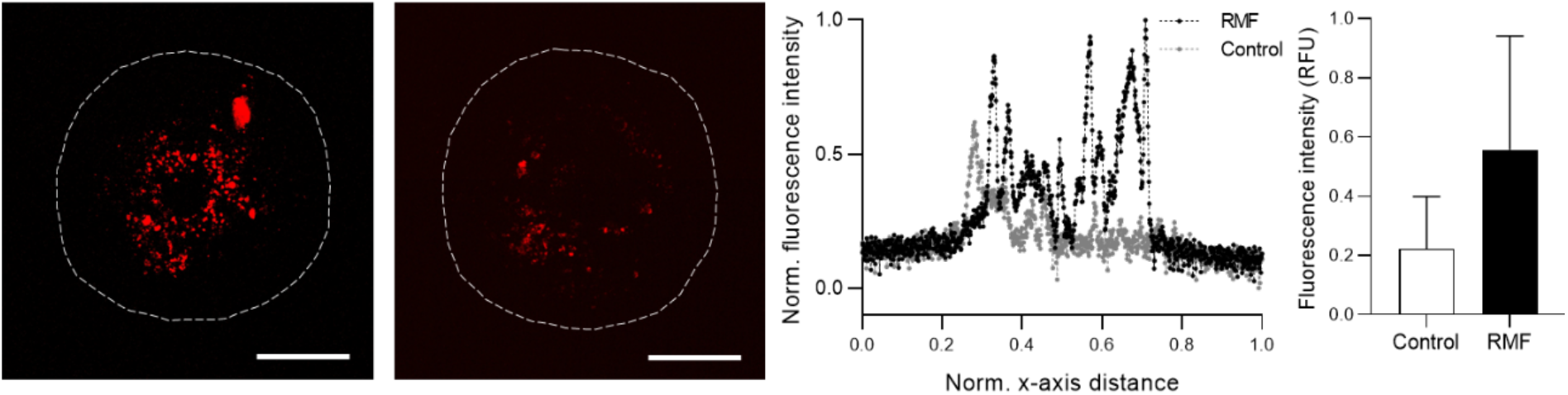
Assessment of MTB colonization in HCT 116 tumor spheroids. A) Representative z-projections from live HCT 116 spheroids following 1 hour exposure to RMF (20 mT and 24 Hz), thorough washing, and incubation for up to 120 h without actuation. MTB were stained with a far-red proliferative stain and images were captured at 120 h. Control refers to unactuated samples. Scale bar = 200 μm. Normalized fluorescence intensity distribution of actuated samples and controls at 120 h (center). Values were normalized to overall minimum and maximum fluorescence intensity values. Spheroid diameter was normalized along x-axis. Image-based quantification of fluorescence intensity values from z-projections at 120 h (right; *n* = 3; mean ± SD).

**Fig. S8.**
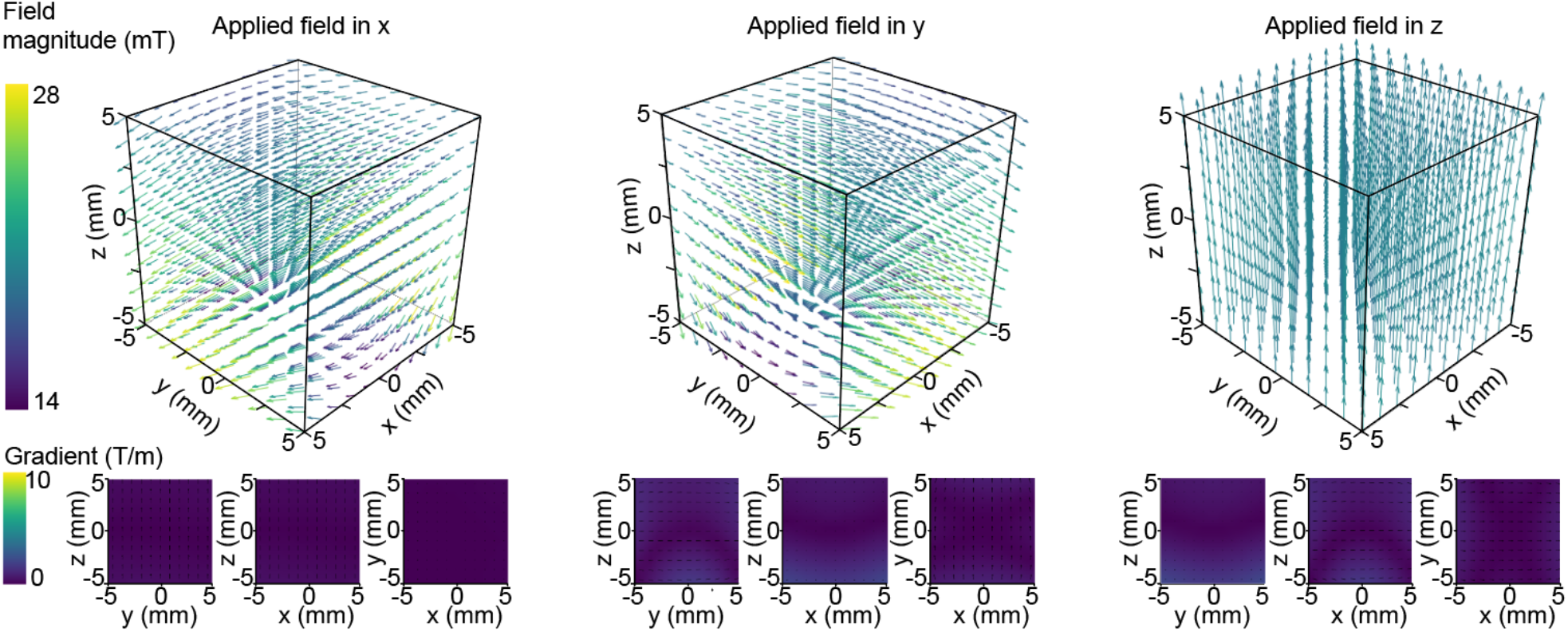
Plots of expected field magnitude and gradients. Fields and gradients produced in the workspace of the magnetic field generator for an applied field of 20 mT in x, y, and z.

**Fig. S9:**
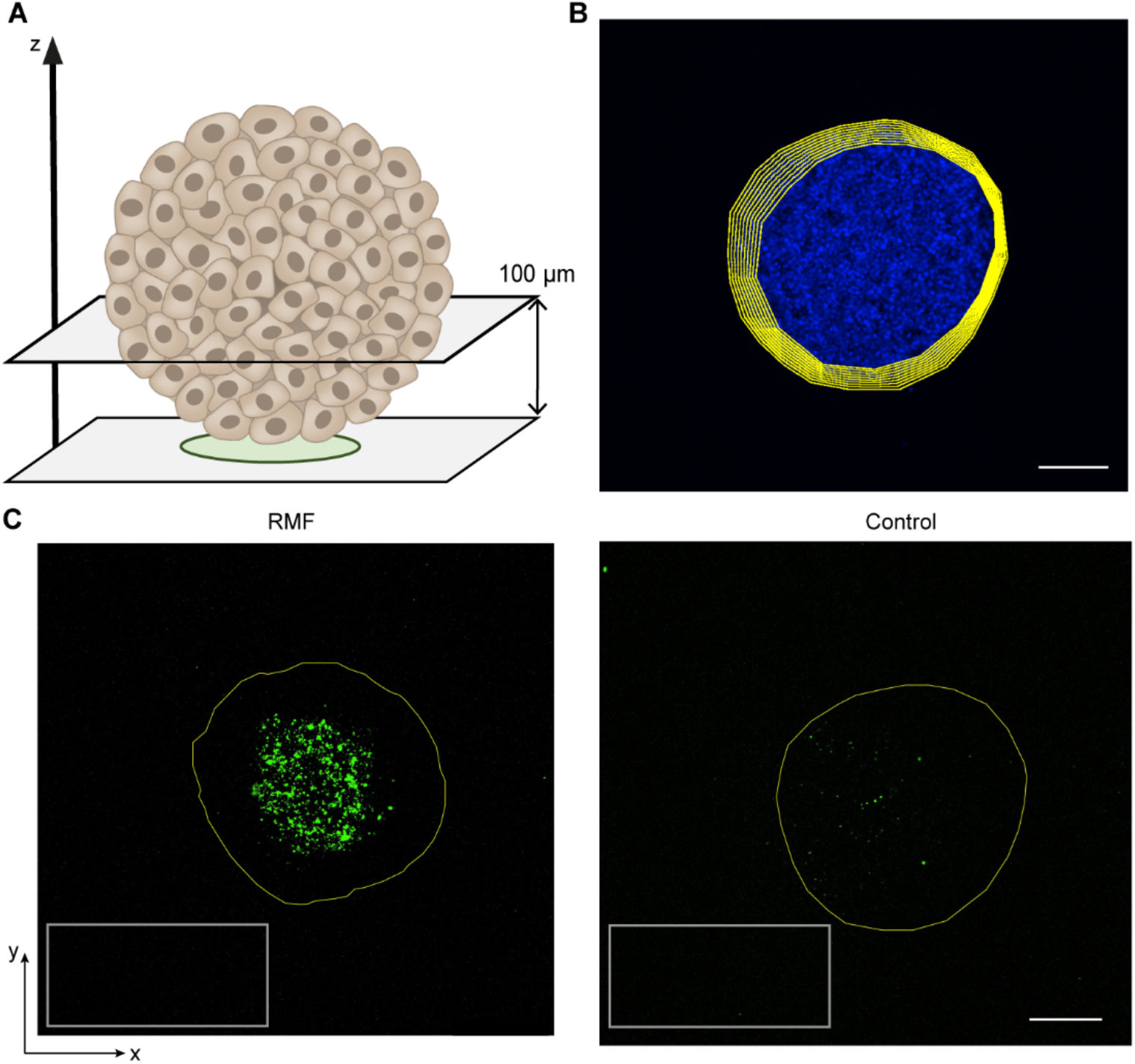
Quantification of fluorescence intensity in spheroids. (**A**) Schematic of MCF-7 spheroid with illustration of z-axis planes. Images were captured at 10 μm intervals from the bottom of the spheroids up to a depth of 100 μm. (**B**) Z-projection of Hoechst image stack with overlaid regions of interest (ROIs). To quantify the average fluorescence of DiO liposomes for each spheroid, ROIs were defined using the Hoechst image stack by tracing a contour around the spheroid at 0 and 100 µm. Intermediate ROIs were defined using interpolation, with approximately 3 µm radial increments. These ROIs were then applied to the corresponding DiO liposome image stack. Scale bar = 200 μm. (**C**) Representative images of sections from DiO liposome image stacks showing an interpolated ROI (yellow line). These fluorescence values were normalized to the average fluorescence of the surrounding media (ROI = gray rectangle). Scale bar = 200 μm.

**Table S1.** Physicochemical characterization of liposomes

| Mean particle diameter [nm] | Mean zeta potential [mV] | Polydispersity index |
| --- | --- | --- |
| $184.2 \pm 3.2$ | $39.8 \pm 0.9 \text{ mV}$ | $0.142 \pm 0.055$ |

## Supplementary Movies

Movie S1: Modeling of liposome diffusion across endothelial monolayer.

Movie S4: Actuation of MTB on HMEC-1 monolayer

Movie S5: MTB actuation into MCF-7 spheroid

